# Purification of filamentous phage ssDNA genomes using Chaotropic salt/silica methods

**DOI:** 10.64898/2026.10.21.760820

**Authors:** Saumel Perez-Rodriguez, Arya Khosravi, Carlos Milla, Paul L Bollyky

## Abstract

Filamentous phages play important roles in molecular biology and bacterial pathogenesis. There is therefore great interest in the genomes of the phages, including M13 and Pf, which infect *Escherichia coli* and *Pseudomonas aeruginosa*, respectively. However, efficient methods for purifying single-stranded DNA (ssDNA) from filamentous phages are scarce and rely mainly on repurposed commercial kits developed for dsDNA. These kits yield variable amounts of ssDNA, varying considerably from one phage to another, their solutions are proprietary and troubleshooting, and customizing the protocol to improve purification is not possible. Here, we developed and optimized two protocols based on ssDNA binding to silica columns in the presence of guanidine hydrochloride and potassium iodide chaotropic salts, for filamentous phages M13KO7 and Pf4. The protocols achieved 60 to 100% recovery, equal to or higher than the Monarch Spin gDNA Extraction and DNeasy Blood and Tissue kits, with similar A_260/280_ and A_260/230_ ratios, intact DNA in agarose gels, and no detectable host genomic DNA by PCR. Recovery depended critically on the chaotropic salts and on ethanol, and did not increase at lower pH, consistent with a water-exclusion model in which hydrophobic interactions and hydrogen bonding dominate over electrostatic repulsion. Phage-specific differences in recovery indicate that optimization may be needed for some phages, which the fully customizable nature of these protocols facilitates. Comparing the virion ssDNA with the double-stranded replicative form (RF) revealed genome variants in both phages, including a shorter M13KO7 form lacking the kanamycin resistance-p15A insert, and illustrates the value of analyzing both RF and ssDNA forms to study filamentous phage biology. Implementing these protocols in routine filamentous phage workflows should facilitate their characterization and engineering.

## Introduction

Filamentous phages belong to the family *Inoviridae* and are characterized by a circular single-stranded DNA (ssDNA) genome encapsidated in a flexible, tube-like particle formed by coat proteins ^1,2^. Unlike classical lytic phages, they do not lyse their host and are therefore generally not used in phage therapy. However, they are widely used in research and biotechnology^3^, and are studied for their role during pathogen infections^4^. *Escherichia coli* M13 and fd filamentous phages are the classical representatives of this group, with M13 being thoroughly characterized and extensively modified^5–7^.

M13 phage has found multiple applications in molecular biology, biotechnology, and materials science. The most important ones have been the development of cloning vectors for molecular biology ^5,8^, the development of multiple display systems for peptides, proteins, single-chain antibodies (scFv) and Fabs fused to different coat proteins in a variety of arrangements for the high-throughput screening of binders and development of therapeutics^9–12^, and the production of ssDNA ^13,14^. Other applications include biosensors, cell imaging probes, immunoassays, drug delivery, tissue regeneration, and targeted cancer therapy^6,7^.

Pf4 phage is other filamentous phage with well-characterized roles in *Pseudomonas aeruginosa* (Pa) biology and microbial pathogenesis ^15^. Pf4 enhances Pa pathogenicity during wound and lung infections by several distinct mechanisms^16,17^. In established biofilms, Pf4 genes are highly expressed, and phage titers increase significantly, leading to the emergence of small-colony variants with enhanced attachment and biofilm development that contribute to Pa virulence *in vivo* ^18,19^. Viral filaments assemble with host- and bacteria-derived polymers to build ordered, liquid-crystal biofilm matrices that help the bacteria withstand desiccation and antibiotics^20^.

When filamentous phages infect the host, the viral positive DNA strand (+) reaches the bacterial cytosol and serves as a template for the synthesis of the complementary negative strand (-) by host RNA and DNA polymerases, producing the double-stranded DNA (dsDNA) genome called the replicative form (RF)^2,8^. Hundreds of RF copies are then synthesized with the help of viral gII complexed with host polymerase and poorly described factors, before the translation of other viral genes is sufficient to start viral strand packaging and virion assembly at the bacterial membrane and phage extrusion into the extracellular medium^1,21^. An intergenic region (IG) contains the viral origin of replication and the packaging signal, where multiple hairpins and sequence-specific elements are necessary for the synthesis of the RNA primer, positive strand extension and ssDNA assembly with viral membrane proteins^22–25^.

M13 phage, specifically, possesses a 6.4 kb genome coding for 11 genes and a ∼500 bp IG sequence between genes IV and II responsible for replication and phage assembly. This viral ori confers DNA replication and production of chimeric M13-like particles when transferred to plasmids in a helper phage-dependent manner^22–24,26^. Given these properties, two kinds of M13-based vectors have been developed, phage vectors and phagemids. An important drawback of phage vectors is, however, the emergence of deletion mutants that outcompete the original population by restoring the enhancer of replication and ejecting the non-viral cloned DNA, resulting in a fast propagation phenotype ^27^. This genetic instability led to the development of phagemids, plasmids incorporating an M13 ori that can be replicated as ssDNA and packaged into phage particles when M13 genes are provided in *trans*. These viral genes can be expressed in *trans* from compatible M13-derived helper plasmids or M13 helper phages ^28–30^. M13KO7 represents a common helper phage used to produce chimeric particles from phagemids, in which a low-copy p15A plasmid origin of replication together with a kanamycin resistance gene (Tn903 kanR) are inserted into the IG. This insertion disrupts the helper phage packaging signal/origin enough that phagemids carrying a wild-type f1/M13 origin are preferentially packaged^31^.

Pf on the other side, integrates as a ∼12.5 kb prophage in the Pa PAO1 strain at the tRNA-Gly site. According to a genetic classification based on genes encoding structural/morphogenesis proteins, Pf4 belongs to a separate lineage and is broadly distributed among clinical Pa isolates^32–34^. Pf4 excision from the prophage is induced by low oxygen, oxidative stress ^15^ and iron-depletion ^35^ growth conditions, mediated by the phage excisionase xisF4, after which it circularizes as a dsDNA RF from which ssDNA is synthesized and viral genes are transcribed to produce viral filaments in the medium^36^.

Given the broad use of M13-derived phages and the well-established role of Pf4 in P. aeruginosa pathogenicity, studying these and other filamentous phages remains crucial. Tracking their mutations by next-generation sequencing (NGS) reveals key information about their properties and their impact on bacterial hosts and mammalian tissues, particularly as genetic instability, including mutations, deletions and rearrangements that favour propagation, has been reported in Pf4 and M13 ^37,38^. Filamentous phages can be sequenced from their double-stranded RF using standard plasmid miniprep methods, but the ssDNA genome is the molecule packaged into virions and transmitted to new hosts, making it the most informative target. RF-based approaches cannot detect changes introduced during ssDNA synthesis, so sequencing virion ssDNA gives a more faithful picture of the phage population actually released into the environment.

Phage gDNA is typically purified by removing free host nucleic acids with DNase/RNase, precipitating virions (e.g., with PEG/NaCl, acetone, TCA or ultracentrifugation), lysing them with proteinase K, detergent and/or chaotropic salts, and capturing the released DNA on silica columns or glass beads, usually with commercial kits (Supplementary Table III)^39–49^. Most of this work has been done on tailed phages with linear dsDNA genomes, ranging from ∼19 kb (Φ29) to ∼500 kb (*Priestia (Bacillus) megaterium* G) ^39,47^. Yield is roughly 0.1–9 µg of DNA of good purity per ml of supernatant, depending on the phage, the titer and the method^43,44,47,48^. However, published protocols also have several limitations. First, they have been optimized and validated on lytic dsDNA phages, so their results are not necessarily valid for ssDNA. Second, many workflows rely on proprietary kits whose composition or lysis conditions are undisclosed ^48–50^, which limits troubleshooting and adaptation to different samples. Finally, those with non-column formats are more laborious, variable and difficult to implement routinely.

In the specific case of filamentous phages, purification methods for their genomic ssDNA are far less developed, covering mainly M13 derivatives ^50–53^ and in one case Pf phages ^54^. These mostly adapt plasmid miniprep or DNA clean-up kits, and were developed mainly to produce ssDNA scaffolds for DNA origami^51,52^ or Pf4 detection^54^. Work on Pf phages was also limited to complex clinical samples, in which phage ssDNA is co-extracted with abundant host dsDNA. Though ssDNA yields of 1–12 µg/ml are comparable to those for dsDNA phages, they are highly phage-dependent. As a result, there is no standardized protocol for obtaining virion ssDNA suitable for sequencing and other downstream applications, whereas RF-based approaches remain the default.

Consequently, we aimed to evaluate whether chaotropic salt/silica methods routinely used to purify dsDNA can be adapted to purify ssDNA from filamentous phages. We selected the filamentous ssDNA phages M13KO7 and Pf4 as models because of their biotechnological and clinical importance and because they differ in host, genome sequence and GC content. We systematically investigated the main chemical and physical factors affecting ssDNA purification, which have not been systematically addressed for ssDNA, to gain insight into the molecular mechanisms driving ssDNA-silica interactions. In addition, we compared the resulting protocols side by side with kit-based protocols commonly used for lytic dsDNA phages, in terms of yield, purity, integrity and performance in downstream applications, to assess their broader applicability to filamentous phages.

## Materials and Methods

### Bacterial cultures and phage propagation

#### Bacterial cultures and identity validation by PCR

Well-validated *E. coli* strains were acquired from commercial vendors while *Pseudomonas* strains were obtained from the Secor lab and validated by molecular techniques^55^. *E. coli* strain NEB5α (New England Biolabs, NEB, C1010S), *Pseudomonas aeruginosa* MPAO1 and its derivatives MPAO1 ΔPf4 and MPAO1 ΔPf4 ΔPf6 were streaked on an LB Miller-agar 1.5% plate and incubated for 16 h at 37°C. MPAO1 ΔPf4 lacks the whole Pf4 prophage locus, while MPAO1 ΔPf4 ΔPf6 lacks both the Pf4 and Pf6 prophages. Liquid cultures were obtained from an isolated colony inoculated in 50 ml of LB Miller medium in a 250 ml Erlenmeyer flask and incubated at 250 rpm and 37°C for 16-18 h. *E. coli* ER2738 strain (Biosearch technologies 60522-1) was streaked on an LB Lennox-agar 1.5% plate supplemented with 20 μg/ml of tetracycline (Tet) and incubated for 16 h at 37°C. An individual colony was inoculated in 50 ml LB Lennox medium supplemented with 20 μg/ml Tet in a 250 ml Erlenmeyer flask and incubated at 250 rpm and 37°C for 16-18 h. All strains were preserved at -80°C in cryovials containing overnight cultures and sterile 40% glycerol at a 1:1 volume ratio.

The three MPAO1 strains were validated by PCR and NGS. Water used throughout the study was nuclease-free, molecular biology-grade. Primers were designed in this study using the Primer3Plus and NCBI Primer-BLAST tools and checked with the IDT OligoAnalyzer web tool. For PCR, 1/OD_600nm_ ml were centrifuged at 8000 x g for 30 min at 4°C, the bacterial pellet was washed twice with 1 ml of sterile phosphate-buffered saline (PBS, 137 mM NaCl, 2.7 mM KCl, 10 mM Na_2_HPO_4_, and 1.8 mM KH_2_PO_4_), resuspended in 100 μl of lysis buffer (100 mM Tris-HCl, pH 8.5, Tween-20 0.05%), incubated at 95°C for 5 min, cooled to room temperature (RT); proteinase K (IBI Scientific IB05406) was added at 240 µg/ml, followed by incubation at 60°C 1400 rpm for one hour and the enzyme was inactivated at 97°C for 15 min. The SN from a 10,000 x g centrifugation for 5 min was stored at -20°C as a template for the respective PCRs.

Standard desalted 25 nmol DNA oligos were ordered from Integrated DNA Technologies (IDT), centrifuged at 15,000 x g for 1 min and resuspended thoroughly by vortexing in water at 100 μM. Ready-to-use primer preparations were then prepared by diluting these 100 μM stocks to a final concentration of 25 μM in the same water. All primers were stored at -20°C. The three MPAO1 strains used in this study were initially validated for the presence of Pf4 and Pf6 prophages using published and newly designed PCRs, adapted to the EconoTaq Plus Green 2x Master Mix (Biosearch technologies 30033-1) and 10-20 μl reactions visualized in 1-2% agarose gels alongside 100 bp (ThermoFisher Scientific SM0243) or 1 kb (NEB N3232S) DNA ladders. PCRs specific to the RF^18^, phage integration left^19^ or right^4^ sites, internal phage sequence^4^ and Pf4-Pa^55^ boundaries were run to assess Pf4 prophage status. Pf6 prophage was identified by a Pf6-Pa boundary PCR ^55^ and the production of its RF (this study). All PCR reactions and running conditions are specified in Supplementary Tables I and II.

For all DNA visualizations, samples and DNA ladders were diluted in water at the required concentration with 6x Gel Loading Dye (NEB B7024S), applied to agarose gels supplemented with 1 μg/ml ethidium bromide and run at a constant 120 V in a Thermo Scientific Owl Easycast B2-BP electrophoresis system. Tris-acetate-EDTA (TAE, 40 mM Tris, 20 mM acetic acid, 1 mM EDTA, pH 7.6) buffer was chosen for gel casting and as the running buffer. Images were acquired in a BioRad Chemidoc XRS system with Image Lab v6.1.0 software and analyzed in ImageJ v1.54g software.

#### Bacterial gDNA purification, quality assessment and sequencing

Bacterial gDNA was purified by the Monarch Spin gDNA Extraction Kit (NEB T3010L) with Cell Lysis Buffer (CLB) following the manufacturer’s simplified protocol for Gram-negative bacteria, which does not require lysozyme. Purified gDNA was quantified using both a Nanodrop 2000 Spectrophotometer with default dsDNA settings and a Quant-iT PicoGreen dsDNA Assay Kit (ThermoScientific P7589) and dsDNA reagent (ThermoScientific P7581) following the manufacturer’s recommendations. DNA quality was evaluated by A_260/280_ and A_260/230_ ratios from Nanodrop measurements and gDNA visualization in a 1% agarose gel alongside a 1 kb DNA ladder. Bacterial genomic DNA was sequenced by Plasmidsaurus using high-depth Oxford Nanopore long-read sequencing with amplification-free library preparation, R10.4.1 flow cells, and Super Accurate basecalling, followed by *de novo* assembly and polishing (Standard service).

Whole-genome alignments were run in the BLAST Ring Image Generator (BRIG v0.95)^56^, while Pf4 and Pf6 loci were specifically compared across 20 kb in Geneious Prime (v2026.1.3) by multiple alignment.

#### Preparation of Pf4 master phage banks

The protocol for the propagation of filamentous Pf4 phage was previously established in the Bollyky lab ^57^. In brief, this starts with the SN of non-infected cultures collected after 18-24 h incubation at 37°C 250 rpm in LB Miller medium. Phage SNs were clarified by centrifugation at 8,000 x g 15 min 4°C and 0.2 μm filtration, precipitated by adding 1/3 volume of cold 16% (w/v) PEG-8000 and 2 M NaCl solution and incubated at 4°C for 24 h. The phage pellet was recovered by centrifugation at 8,000 x g 4°C 30 min and carefully solubilized in saline-magnesium (SM) buffer (Tris-HCl 50 mM, NaCl 100 mM, MgSO_4_ 8 mM) supplemented with 0.01% gelatin^58^ at 1/40 SN volume. Phages were then dialyzed three times against 200x volumes of the same buffer, 0.2 μm sterile filtered and stored at 4°C. Pf4 quantification was done by classical plaque assay in Petri dishes made from pre-warmed (50-55°C) LB Miller-agar 0.5% with 400 μl of a mid-log phase MPAO1 ΔPf4 culture (OD_600nm_∼0.5-0.6). Ten-fold dilutions were made in PBS and 10 μl were spotted in Petri dishes, which were then incubated at 37°C for 14-16 h. Further propagation rounds involved cyclic superinfection of MPAO1 cultures at OD_600nm_=0.3-0.4 with a multiplicity of infection (MOI) ∼50, 2 h incubation at 37°C 250 rpm, 100-fold dilution in fresh LB Miller medium and another 14-16 h incubation under the same conditions.

#### Preparation and quantification of working Pf4 batches

For the preparation of three biological replicates for this study, one batch of the Pf4 master phage bank (1.8 x 10^10^ plaque forming units (PFU)/ml) was 10-fold diluted in sterile Tris-buffered saline (TBS, 50 mM Tris-HCl, pH 7.5, 150 mM NaCl) and 100 μl of each of the 10^6^-10^7^ dilutions was mixed with pre-warmed (50-55°C) 20 ml LB Miller-agar 0.5% and 400 μl of an overnight culture of MPAO1 ΔPf4 ΔPf6 strain. The medium was poured into Petri plates, dried in the laminar flow hood until solid and incubated at 37°C for 14-16 h. Three individual plaques were inoculated in 50 ml of LB Miller medium, supplemented with overnight MPAO1 ΔPf4 ΔPf6 cultures diluted at an OD_600nm_=0.05 and incubated at 37°C 250 rpm for 24 h. SNs were clarified by centrifugation at 8,000 x g 15 min 4°C and 0.2 μm filtration and stored at 4°C. Phage quantification was done by the same plaque assay developed for Pf4 master phage banks.

#### Propagation and quantification of phage M13KO7

The kanamycin-resistant *E. coli* helper phage M13KO7 (NEB N0315S) was propagated in the ER2738 strain in LB Lennox medium supplemented with 20 μg/ml Tet. To that end, a 50 ml fresh culture was prepared in a 250 ml Erlenmeyer flask by diluting an overnight bacterial culture to a final OD_600nm_=0.05 and infected with a 1000-fold dilution of the commercial phage preparation (1.0 x 10_11_ PFU/ml), representing a MOI of about 2.5. The culture was incubated at 37°C 250 rpm for 90 min, after which kanamycin was added at 70 μg/ml and incubation was resumed for another 14-18 h. SN was recovered by centrifugation at 6,000 x g 15 min 4°C and 0.2 μm filter-sterilized and stored at 4°C. The M13KO7 master phage banks were prepared after phage precipitation with 1/5 SN volume of 20% PEG-8000 and 2.5 M NaCl, 4°C incubation for 24 h, centrifugation at 5,000 x g 45 min 4°C, pellet solubilization in 1/20 SN volume of TBS, three times dialysis against 200x volumes of TBS at 4°C, 0.2 μm sterile filtration, and storage at 4°C. M13 phage quantification was done by the classical plaque assay with some modifications. A fresh ER2738 culture was initiated at OD_600nm_=0.05 and incubated at 37°C 250 rpm until mid-log phase (OD_600nm_∼0.5). Agar plates were prepared by mixing 20 ml of pre-warmed (50-55°C in a water bath) LB Lennox-agar 0.5% with 400 μl of ER2738 culture and Tet at 20 μg/ml, poured into a Petri dish and left open until the medium solidified. M13KO7 SN was 10-fold diluted in sterile TBS in a cell culture 96-well plate, and 10 μl of each dilution was spotted on the plate and dried in the laminar flow hood. Plates were then incubated at 37°C for 14-16 h, individual plaques were counted and the phage titer was calculated as PFU/ml at the corresponding dilution. Three biological replicates were obtained.

### Evaluation of ssDNA purification methods and influencing key factors

#### Pretreatment of phage-enriched SNs prior to ssDNA purification

DNase I (Roche 10104159001) was dissolved in 10 ml of autoclaved 20 mM Tris-HCl, 2 mM MgSO_4_, 2 mM CaCl_2_, pH 7.5 solution following the manufacturer’s recommendations, and later diluted in the same volume of molecular biology-grade glycerol, aliquoted and stored at -20°C (5 mg/ml, 10 U/μl stock). RNase A (BioBasic RB0473) was dissolved in 20 mM Tris-HCl, 30 mM NaCl, pH 7.5 at 40 mg/ml, and later diluted, aliquoted and stored the same way (20 mg/ml stock).

One ml of phage-enriched and clarified SN at room temperature (RT) was processed per silica column. Contaminating DNA and RNA, not encapsulated in phages, were digested in the presence of MgSO_4_ 10 mM, CaCl_2_ 1 mM, RNase A 50 μg/ml and DNase I 20 U/ml, for 1 h at 37°C. Nucleases were then inhibited by EDTA 20 mM. Next, phages were precipitated with NaCl 0.5 M and cold PEG-8000 10% Precipitates were stored at 4°C for 18 h, centrifuged at 15,000 x g 30 min 4°C, SN removed and the phage pellet used for ssDNA purification according to each method.

#### Monarch Spin gDNA Extraction Kit

A kit optimized for the isolation of genomic DNA from a broad range of sample types, such as mammalian cells and tissues, bacteria and yeast, was initially tested for its performance on the small Pf4 phage ssDNA genome before testing the broader DNeasy Blood & Tissue Kit (Qiagen 69504), which covers all these and many other non-standard samples. The phage pellet was resuspended in 100 μl of PBS and proteinase K was added at 4 μg/μl. A hundred microliters of Cell Lysis Buffer (CLB) were next added, well mixed and the sample incubated for 2 h at 56°C and 1400 rpm in a ThermoMixer C. Samples were brought to RT for 10 min, briefly spun down and mixed thoroughly with 400 μl of Binding Buffer. The samples were transferred to a gDNA Purification column pre-inserted into a collection tube and incubated for 10 min at RT. The column was centrifuged first for 3 min at 1,000 x g, followed by 1 min at 12,000 x g, and washed three times with 500 μl of Wash Buffer for 1 min at 12,000 x g. Another centrifugation step was done to eliminate residual buffer from the column, and 100 μl of preheated elution buffer (10 mM Tris-HCl, 0.1 mM EDTA, pH 9.0, 60°C) was added to the column and incubated for 10 min at RT. DNA was eluted by centrifugation for 2 min at 12,000 x g, and the eluate was applied for a second time to the column and centrifuged under the same conditions. New collection tubes were always used in between centrifugation steps.

#### ssDNA purification by DNeasy Blood & Tissue kit

ssDNA was purified from PEG/NaCl-precipitated phage pellets using the DNeasy Blood & Tissue Kit (Qiagen 69504). The pellet was resuspended in 200 μl of PBS and viral particles were disassembled with the addition of 2 μg/μl of proteinase K, 200 μl (1:1 volume ratio) of AL buffer and sample incubation for 2 h at 56°C under 1400 rpm agitation in a ThermoMixer C. The tubes were subsequently pulse-centrifuged, thoroughly mixed with 200 μl (1 volume) of pure molecular biology-grade ethanol and transferred to a DNeasy Mini spin column. ssDNA binding was performed first at 1000 x g for 3 min, followed by 1 min at 12,000 x g. The column was washed for 1 min at 12,000 x g with 500 μl of AW1 buffer followed by 500 μl of AW2. A second AW2 wash was done for 3 min at 17,000 x g. All collection tubes were changed after each centrifugation step. ssDNA was finally eluted with 100 μl elution buffer (Tris-HCl 10 mM pH 8.5) pre-warmed to 60°C, by centrifugation for 2 min at 12,000 x g, and the eluate was passed a second time through the column to increase recovery. Purified phage DNA was stored at -20°C for all further analyses.

#### ssDNA purification by chaotropic salts method

A chaotropic-based ssDNA purification protocol was fully customized from PEG/NaCl-precipitated phage pellets. These pellets were resuspended in 200 μl of PBS supplemented with Tween-20 0.5% and Triton X-100 1%, and then mixed with an equal volume of 3 M chaotropic salt (guanidine hydrochloride (GuHCl) or potassium iodide (KI)) and 20 μl of 20 mg/ml proteinase K. Samples were incubated for 2 h at 56°C under 1400 rpm agitation in a ThermoMixer C. Tubes were briefly spun down, 400 μl of pure ethanol (2 volumes) was added and mixed thoroughly by vortexing. Samples were then transferred to a silica column, incubated for 10 min, and centrifuged at 1000 x g for 3 min, followed by another 2 min at 12,000 x g. The column was extensively washed three times by incubating it for 5 min with 500 μl of washing buffer (1 mM Tris-HCl, 10 mM NaCl, pH 7.5, ethanol 80%) and centrifugation for 2 min at 12,000 x g. Two more washes were performed using the same incubation and centrifugation settings with 500 μl of pure ethanol. An additional centrifugation of the empty column removed any remaining buffer prior to DNA elution. A hundred microliters of warm elution buffer (Tris-HCl 10 mM pH 8.5, 60°C) was added to the column, incubated for 10 min and centrifuged for 2 min at 12,000 x g. This eluate was passed again through the column to increase recovery, and the purified phage DNA was stored at -20°C for all further analyses. All collection tubes were changed between centrifugations.

#### Evaluation of impact of chemical and physical factors during ssDNA purification by the chaotropic salts method

Different chemical and physical factors involved in ssDNA purification were evaluated for their impact on DNA recovery and quality from Pf4 phage SNs. Silica columns were compared across three common column formats, EconoSpin All-In-One DNA Only Mini Spin Column (Epoch Life Science 1920-250), QIAprep 2.0 Spin Miniprep Column (Qiagen 27115) and EZ-10 Spin Column (BioBasic BS413), using the GuHCl and KI purification protocols. The other chemical factors tested on the Econo DNA columns using the standard KI purification protocol were the presence/absence of the KI chaotropic salt and its timing (before/after incubation at 56°C), of the detergents in the phage-dissolving PBS buffer and of proteinase K, and ethanol concentration after chaotropic salt addition. The initial amount of ssDNA available for purification was additionally correlated (Pearson correlation) with, and used to predict (linear regression), ssDNA recovery by precipitating 0.125 – 1 ml volumes of clarified phage SN and proceeding to KI-mediated ssDNA purification of the respective phage pellets. The physical factors, heat, agitation and incubation time, were evaluated by skipping the heat and agitation treatment during incubation or proceeding to DNA binding to the silica column without any incubation or sample treatment (all three factors together).

#### Quantification of purified phage ssDNA and quality assessment

Purified ssDNA was quantified using both a Nanodrop 2000 Spectrophotometer with default ssDNA settings and a Qubit ssDNA Assay Kit (Thermo Scientific Q10212) following the manufacturer’s recommendations. ssDNA quality was assessed by the A_260/280_ and A_260/230_ ratios from Nanodrop measurements.

#### Detection of gDNA host contamination in purified phage ssDNA

The presence of contaminating host gDNA in purified phage ssDNA was validated by PCR to confirm NGS results. MPAO1 gDNA in Pf4 genomes was tested using the same Pf6 multiplex assay used previously to identify Pf6 prophage in MPAO1 strains ^55^. For *E. coli* gDNA contamination, a PCR targeting the characteristic structure of the O-antigen biosynthesis (rfb) locus in K-12 strains was adopted ^59^. All reaction and PCR running conditions are detailed in Supplementary Tables I and II.

### Validation of downstream applications for purified phage ssDNA

#### NGS of phage genome

ssDNA was sequenced as a plasmid by Plasmidsaurus using an amplification-free Oxford Nanopore long-read sequencing workflow. Libraries were prepared with v14 chemistry from sequence-independently linearized plasmids and sequenced on a PromethION P24 instrument using R10.4.1 flow cells to generate a high-accuracy consensus sequence. DNA analyses and alignments were done in SnapGene v8.2.3 software.

#### Phage identity by genome restriction patterns and PCR

Besides DNA sequencing, restriction patterns and PCR of purified phage genomes can be used as quick and inexpensive methods to evaluate phage identity. Pf4 ssDNA restriction patterns were obtained from *KpnI* (NEB R3142S) and *EcoRI* (NEB R3101S) enzymes, with one and four restriction sites in this phage genome, respectively. Given that many restriction enzymes have shown a lower activity on ssDNA substrates, *KpnI* and *EcoRI* were adjusted from 20 U/μl as reported by the manufacturer to 12 U/μl (60% residual activity) and 8 U/μl (40% residual activity), respectively, according to REBASE^60,61^. Hence, a larger enzyme volume was used to deliver 20 U of effective activity. Phage DNA (1 μg) was mixed with 20 U of effective enzyme activity, 1x CutSmart buffer and water in a 50 μl reaction and incubated for 2 h at 37°C. Two hundred nanograms were mixed with 6x loading buffer and separated in a 1% agarose gel alongside a 1 kb DNA ladder.

Enzymes *NdeI* (NEB R0111S) and *SmaI* (NEB R0141S) with three and two sites in the M13KO7 genome, respectively, were chosen to obtain restriction patterns for this phage. *NdeI* activity was adjusted from 20 U/μl to 14 U/μl (70% residual activity), while *SmaI* was used as provided by the manufacturer given the high enzyme concentration that would otherwise be needed based on its 10% residual activity on ssDNA. Five hundred nanograms of ssDNA were incubated with 10 U of each enzyme, 1x CutSmart buffer and water in a 25 μl reaction for 2 h at 37°C, mixed with 6x loading buffer and applied to a 1% agarose gel.

RFs are circular dsDNA phage genomes that are replicated in the bacterial cytosol and serve as a template for gene transcription and translation of phage proteins and from which the positive strand is also amplified and subsequently packaged into extruding virions^2^. Given their plasmid-like behavior, RFs were isolated from the same infected bacteria used for phage propagation by pelleting 5 ml cultures at 8,000 x g 15 min 4°C and purifying the intracellular dsDNA following QIAprep Spin Miniprep kit’s instructions. Purified phage RF was quantified in a Nanodrop 2000 Spectrophotometer using default dsDNA settings and sequenced by Plasmidsaurus using the standard plasmid service. Restriction patterns from the RF were obtained with the original enzyme activity provided by the manufacturer and under the same reaction conditions as ssDNA.

ssDNA identity from Pf4 was confirmed by PCR directed against the gII-PA0717 region, while for M13KO7 an internal gIII region was chosen. As M13 gIII fusion constructs are made in the N- or C-terminal ends, this gIII PCR can be used as a universal M13 identity test for phages and phagemids. All reaction and running conditions have been disclosed in Supplementary Tables I and II.

#### Prediction of ssDNA digestion based on local DNA structure

Residual activity of each enzyme on ssDNA was obtained from the oligonucleotide substrates listed in REBASE ^60^. Because intermolecular pairing could render a recognition site double-stranded, homodimer formation of the REBASE oligonucleotides was calculated with the DINAMelt server ^62^, using the two-state melting (hybridization) application with the same sequence entered for both strands. Calculations were performed for DNA at 37 °C with [Na⁺] = 50 mM and [Mg²⁺] = 10 mM, approximating the ionic conditions of the rCutSmart buffer used for phage ssDNA digestions. All other parameters were left at default values. For each oligonucleotide, the free energy (ΔG), melting temperature (Tm) and predicted base pairing of the duplex were recorded, and the oligonucleotide was considered able to form a homodimer when the melting temperature (T_m_) exceeded 37°C and the recognition site was located within the paired region.

Secondary structure of phage ssDNA was calculated in SnapGene v8.2.3 using ViennaRNA: RNAfold for minimum free energy structures and partition-function base-pair probabilities, and RNAsubopt for suboptimal structures within 5 kcal/mol of the minimum^63^. Calculations used the DNA energy parameters at 37°C and the default ViennaRNA salt conditions ^64^. For the phage ssDNA, a window of 100 nt upstream and downstream of each restriction site was analyzed. A recognition site was considered paired when all its nucleotides were base-paired in the minimum free energy (MFE) structure and had base-pair probabilities ≥70%. Sites were also checked in the suboptimal structures within 5 kcal/mol of the minimum, and a site was reported as paired only if this classification held in >50% of them.

### Estimation of ssDNA purification efficiency and major molecular mechanisms involved

#### Quantification of total phage ssDNA in phage-enriched SNs

The phage pellet was precipitated from 1 ml of clarified phage SN following the pretreatment of phage-enriched SNs described above and resuspended in 100 μl of PBS. The sample was incubated at 100°C for 15 min, brought to RT and briefly spun down. Quantification was done with the Qubit ssDNA Assay Kit on 10-fold and 20-fold PBS-diluted samples, following the manufacturer’s instructions.

#### Decreasing repulsive electrostatic forces between ssDNA and silica by chaotropic salt-mediated ssDNA purification under acid conditions

ssDNA purification protocols based on chaotropic salts made use of PBS as the buffer for sample preparation, and of a high ethanol concentration that dehydrates ssDNA and favors hydrophobic interactions with silica. However, it has been reported that an acidic pH could favor DNA binding to silica by decreasing the negative charge on the silica surface and the electrostatic repulsion with the negatively charged phosphate groups in the DNA, while increasing the hydrogen bonding^65^. To evaluate the relative contribution of electrostatic forces to DNA-silica interactions, a purification protocol under acidic conditions was developed and compared with PBS-based results. To that end, the PBS phage-dissolving buffer for the pellet was replaced with 100 μl of 0.2 M acetate buffer pH 5.0 supplemented with NaCl 0.2 M, Tween-20 0.5% and Triton X-100 1.0%, while keeping the rest of the protocol the same. pH variations across these protocols were measured at the steps of pellet solubilization (buffer used for dissolving the phage pellet after PEG/NaCl precipitation), phage disaggregation (solution containing the solubilizing buffer plus the chaotropic salt), ssDNA binding to column (solution containing the solubilizing buffer, chaotropic salt and ethanol) and ssDNA elution (elution buffer) in the respective solutions without ssDNA at the ratios and order established in the protocols.

## Statistical analysis

All quantitative experiments were performed with three independent biological replicates (independently propagated phage batches, n=3). Individual values and means ± standard deviation (SD) are shown in the figures. Data analyses and graphical representations were performed in GraphPad Prism v11.1.0. Comparisons among three or more groups were made by one-way ANOVA followed by Tukey’s multiple comparisons test, and comparisons between two groups by unpaired two-tailed t test with Welch’s correction. The relationship between initial ssDNA availability (SN volume, ml) and DNA recovery was evaluated by Pearson correlation and simple linear regression (all individual replicates, n=5). Statistical significance was set at p<0.05, and exact p-values are reported unless indicated otherwise. Relative abundance of DNA fragments in agarose gels was quantified in ImageJ v1.54g. No data were excluded from the analyses.

## Results

### Chaotropic-based purification protocols are effective for isolation of Pf4 phage ssDNA

Before phage propagation, all the Pa strains were verified for the presence/absence of the prophages Pf4 and Pf6 in their genomes by PCR (**Supplementary Figure 1 and Supplementary Tables I-II**) and NGS (**Supplementary Figure 2**). Pf4 phage was then propagated from individual plaques in MPAO1 ΔPf4 ΔPf6 infected cells in liquid medium for further purification of its ssDNA genome (**Supplementary Figure 3**). Pf4 phage was amplified to 5.3 ± 3.2 x 10_10_ PFU/ml, quantified in MPAO1 ΔPf4 ΔPf6 lawns by the classical plaque assay, a titer in line with reported values for the release of super-infective Pf4 phages from MPAO1 biofilms^37,66^.

After successful Pf4 production and in agreement with the common workflow for phage DNA purification, a sample preprocessing step comprising SN clarification by centrifugation and 0.2 μm filtration, nuclease treatment to degrade non-encapsulated host nucleic acids, and PEG/NaCl phage precipitation was done (**Figure 1a**). Phage ssDNA was then purified in consecutive stages: phage solubilization and disaggregation to release the viral genomes, sample conditioning and binding to a silica column, removal of contaminant proteins and salts by several washes with ethanol in low-salt buffer, and DNA elution with a warm, low-salt, high-pH buffer (**Figure 1b**). A_260/280_ and A_260/230_ ratios were used to assess contamination with proteins and A_230_-absorbing chemicals, respectively, and, together with agarose gels, gave an overall assessment of purity and integrity of the purified phage genome.

**Figure 1.**
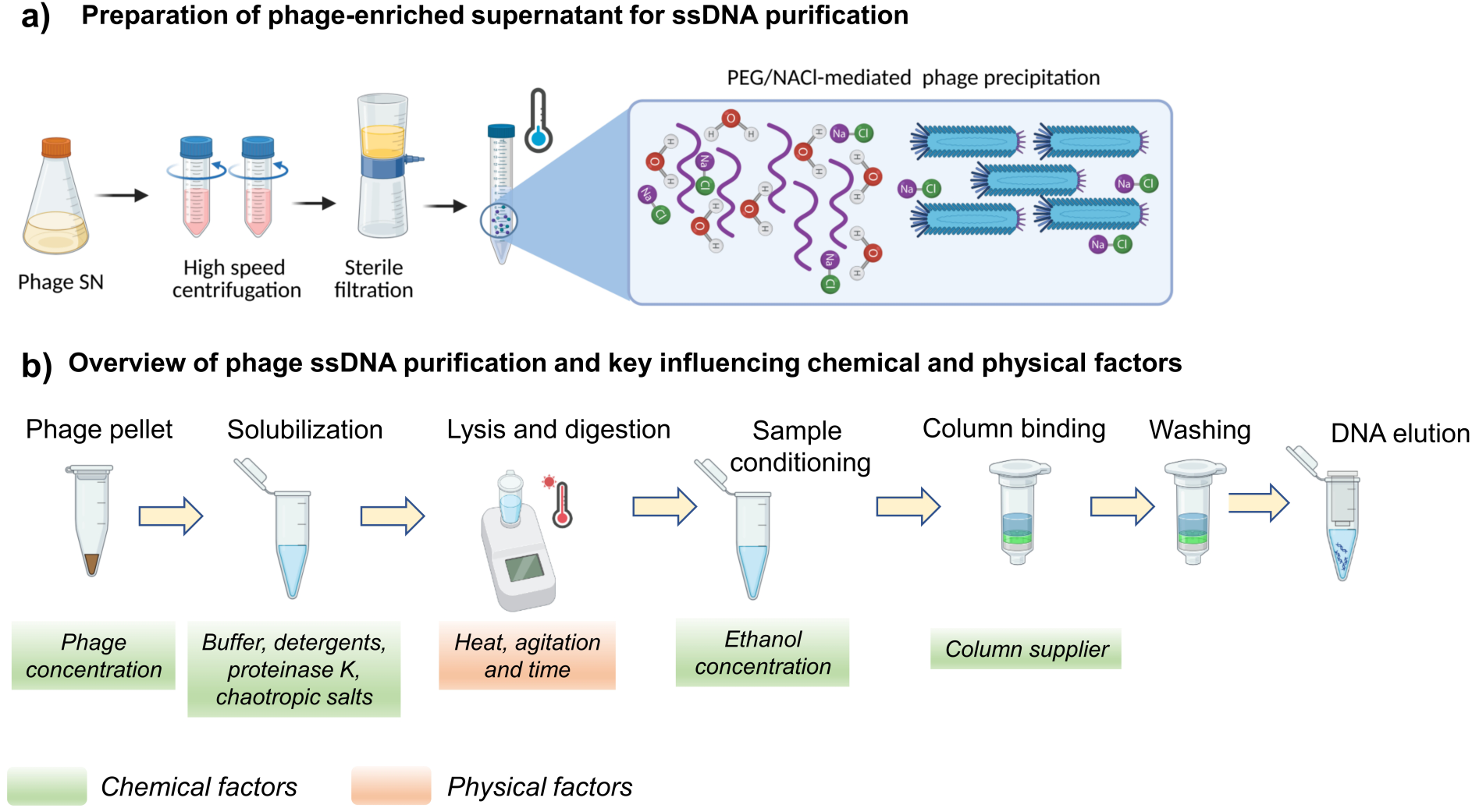
Overview of the ssDNA purification process and factors involved. **a)** Pretreatment of phage-enriched supernatants (SN) before ssDNA purification, involving medium clarification by centrifugation and 0.2 μm filtration, a nuclease digestion step (DNase/RNase) to remove non-encapsidated nucleic acids followed by polyethylene glycol (PEG)/salt-mediated phage precipitation. **b)** General workflow for ssDNA purification with silica columns, divided into consecutive steps: phage solubilization, ssDNA binding to the column, washing and elution. Chemical and physical factors involved in the process were evaluated for their impact on DNA recovery and quality. Created in BioRender. Perez Rodriguez, S. (2027) https://BioRender.com/ldmn42o.

Among the commercial kits attempted for phage DNA purification, those developed for genomic DNA have been used successfully with lytic phages from *E. coli*, *Bacillus subtilis*, *Priestia (Bacillus) megaterium* and *Xanthomonas oryzae* (**Supplementary Table III**) ^44,47^. In line with these precedents, purification of the Pf4 phage genome was first attempted with the Monarch Spin gDNA Extraction Kit, giving a poor recovery of 1.66 ± 0.14 μg/ml SN, although maintaining an acceptable quality (A_260/280_ of 1.88 ± 0.01, and A_260/230_ of 1.53 ± 0.06) and a single pure ssDNA band on 1% agarose gel (**Figure 2a**). While these kits extract and purify genomic DNA from a broad range of samples, they have been tailored to high-molecular-weight (HMW) dsDNA, which probably explains the effective binding of linear 64-500 kb phage dsDNA and the low recovery of the small circular ssDNA from Pf4.

**Figure 2.**
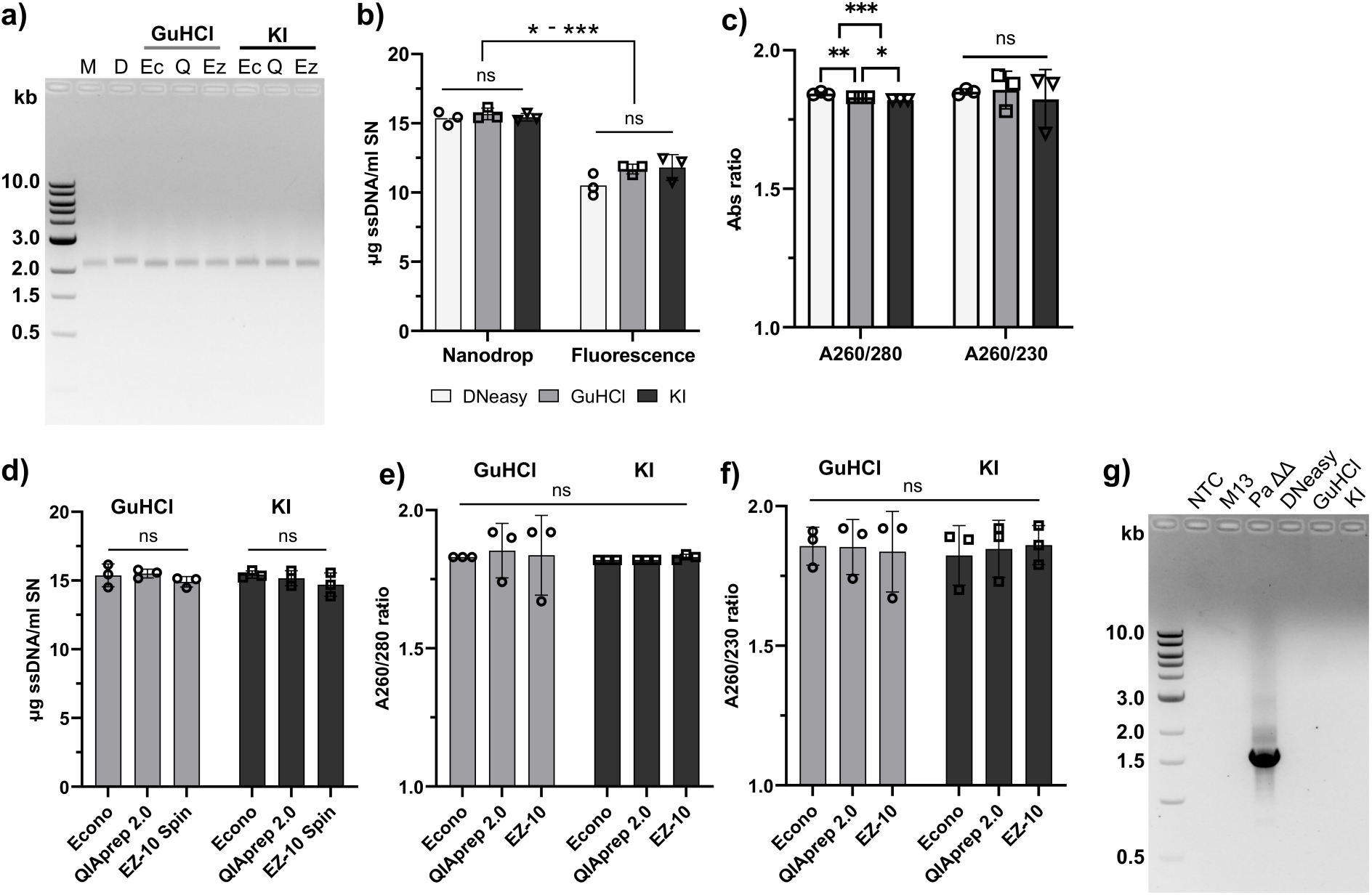
Comparison of Pf4 ssDNA recovery and quality between commercial kits and chaotropic-based protocols. **a)** Evaluation of ssDNA integrity and purity of all purification protocols on a 1% agarose gel. 200 ng of total DNA was applied per lane alongside a 1 kb DNA ladder. M: Monarch Spin gDNA Extraction Kit, D: DNeasy Blood & Tissue Kit, Ec: EconoSpin All-In-One DNA Only Mini Spin Column, Q: QIAprep 2.0 Spin Miniprep Column, Ez: EZ-10 Spin Column. **b-c)** ssDNA recovery and quality control (A_260/280_ and A_260/230_ ratios) of ssDNA purified with the standardized protocols using DNeasy or EconoSpin silica columns, as indicated. SN: phage-enriched supernatant. **d-f)** Performance of three silica column formats (Econo, QIAprep 2.0 and EZ-10 Spin) in GuHCl- and KI-based DNA purifications, in terms of recovery and quality. **g)** Detection of contaminating MPAO1 host gDNA in Pf4 ssDNA purified with the DNeasy, GuHCl and KI protocols, by PCR targeting the Pf6 prophage locus. NTC: non-template control; M13: PEG-precipitated M13KO7 phage, Pa ΔΔ: lysate from *Pseudomonas aeruginosa* (Pa) MPAO1 ΔPf4 ΔPf6 strain. Data are presented as mean ± SD of three independent biological replicates (n=3). Statistical significance was assessed by one-way ANOVA followed by Tukey’s post hoc test for comparisons among three or more groups, and by unpaired t test with Welch’s correction for comparisons between two groups. ns: not significant, *: p<0.05, **: p<0.01, ***: p<0.001.

Since chaotropic-based approaches have been reported for dsDNA phages infecting *Streptococcus uberis*, *Staphylococcus aureus*, *Salmonella sp.*, *E. coli*, *Enterococcus faecalis* and *Geobacillus stearothermophilus* ^40,43,46^, we decided to proceed with these purification methods to capture the total DNA present in samples, independently of DNA nature and size. The commercial DNeasy kit and customizable in-house protocols using the chaotropic salts GuHCl and KI were compared side by side in terms of recovery, quality and purity.

A first evaluation on Econo silica columns showed that chaotropic-based protocols recovered similar Pf4 DNA amounts from SN to the DNeasy kit (**Figure 2b**, Nanodrop p=0.6459, Fluorescence p=0.1364), regardless of the quantification method used, all being protein-free (A_260/280_ ratio > 1.8) with no differences in A_260/230_ ratios (p=0.8450, **Figure 2c**). As expected, Nanodrop overestimated ssDNA significatively by a factor of 1.37 ± 0.08 across all methods (**Figure 2b**, DNeasy p=0.0019, GuHCl p=0.0003, KI p=0.0158), as previously reported for dsDNA from other sources ^67^, and similar to the ∼2-fold overestimation reported for prokaryotic gDNA^68^. Throughout this study, significant differences between samples were detected with both quantification methods, and spectrophotometry consistently overestimated yields relative to fluorescence. The chaotropic-based protocols showed comparably high recovery (**Figure 2d**, GuHCl p= 0.4342, KI p= 0.3842) and purity (**Figure 2e-f**, A_260/280_ p=0.9922, A_260/230_ p=0.9974) with two other commercial silica columns for plasmids, QIAprep 2.0 Spin Miniprep and EZ-10 Spin columns, indicating that the silica source does not play a critical role in the process. All commercial and chaotropic-based protocols showed a pure band by agarose electrophoresis (**Figure 2a**) and no evidence of contamination with host gDNA, as assessed by a multiplex Pa-specific PCR against the Pf6 locus and boundaries (**Figure 2g**). All protocols yielded high-quality ssDNA, although with different efficiencies. The chaotropic-based protocols showed recovery comparable to the DNeasy kit and significantly higher than the Monarch Spin gDNA Extraction Kit.

### Chaotropic-based protocol performance is shaped by ethanol and salt concentrations and correlates with ssDNA input

Several chemical and physical factors are involved in silica-mediated DNA purification (**Figure 1b**), with different relative impacts on purification parameters. Factors like proteinase K ^69^, chaotropic salt nature and concentration^70–72^, input DNA^72^ and ionic strength^73^ have been shown to modulate the recovery of dsDNA, while others like surfactants for sample solubilization ^69^, temperature and incubation time of DNA-silica ^69,73^, and incubation time and temperature during elution^73^ have shown to not change the output. In line with this previous evidence for dsDNA, we evaluated the effect of several parameters on ssDNA recovery and purity using the Pf4 phage and the KI-based protocol as a model.

Decreasing the ethanol concentration in solubilized phage genome samples significantly reduced the genome recovery (**Figure 3a**, Econo, QIAprep 2.0 and EZ-10 columns p=0.0006, p=0.0356 and p=0.0022, respectively) and quality (**Figure 3c**, A_260/230_ ratio for Econo, QIAprep 2.0 and EZ-10 columns, p=0.0423, p=0.0287 and p=0.0162, respectively) while keeping the preparation protein-free (**Figure 3b**, A_260/280_ ratio >1.8 in all cases), on all the three silica columns used before. The rest of the evaluated factors, all tested on Econo silica columns, also gave protein-free preparations (**Figure 3e**, A_260/280_ ratio > 1.8 in all cases). Among them, only KI concentration (**Figure 3d-f**) and DNA input (**Figure 3g**) significantly modified recovery (DNA yield p-values for KI concentration p=0.0182 and DNA input p<0.0001) and DNA quality (A_260/230_ ratio p-values for KI concentration p=0.0087 and DNA input p<0.0001). KI chaotropic salt was necessary for maximum recovery and purity, while its addition before or after sample incubation under heat and agitation was irrelevant (p>0.9999). Although the factors, detergents and proteinase K for sample solubilization, and agitation, heat and incubation time, did not modify the process yield (p >0.9999, p=0.6999 and p=0.9542, respectively) and quality (A_260/230_ p=0.4016, p=0.9442 and p=0.9955, respectively), the absence of proteinase K and triple agitation/heat/incubation introduced high variability in recovery, with a coefficient of variation (CV) of 32-35% versus less than 1% in control samples, supporting their use to maintain reproducibility. DNA input, varied by processing 0.125-1 ml of SN, was the third parameter with a meaningful impact, showing a strong positive correlation with recovery (**Figure 3g**, p<0.0001, R^2^=0.997) and purity (**Figure 3h**, A_260/230_ ratio, p=0.0266, R^2^=0.847). These relationships were well described by linear regression. Recovery increased linearly with input (**Figure 3g**; Y = 14.63X + 0.49, R² = 0.990, p<0.0001, n=5), with an intercept not significantly different from zero. The A_260/230_ ratio also increased with input, although with a poorer fit (**Figure 3h**; Y = 0.93X + 0.98, R² = 0.799, p<0.0001, n=5). The observation that ethanol concentration, chaotropic salt, and DNA input are the primary determinants of purification performance paves the way for customization and optimization of this protocol for challenging samples, an approach that is difficult to apply to commercial kits based on proprietary solutions.

**Figure 3.**
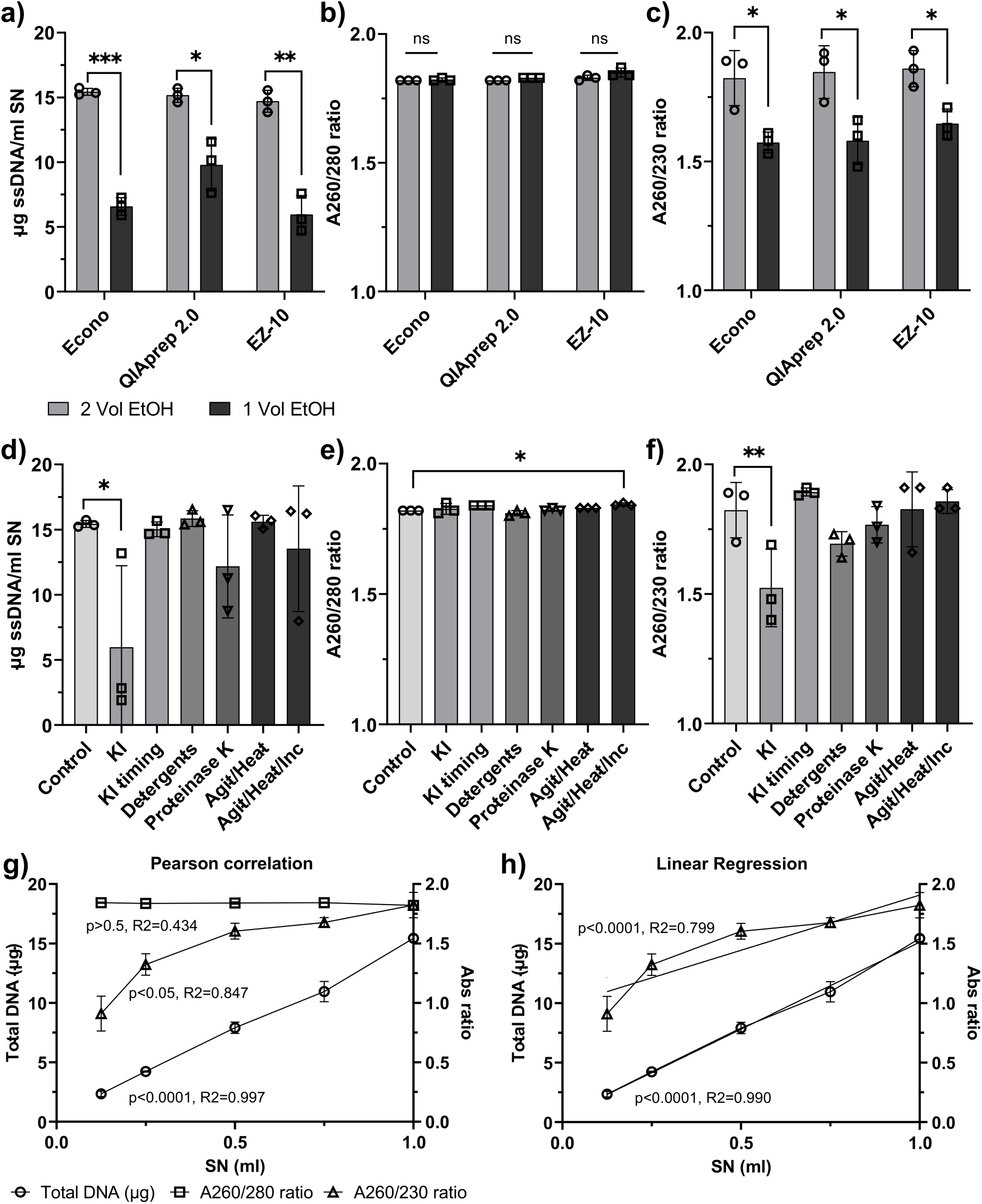
Effect of chemical and physical factors on Pf4 ssDNA recovery and quality for the KI-based protocol. **a-c)** Impact of ethanol concentration on ssDNA recovery and quality control (A_260/280_ and A_260/230_ ratios) of purified ssDNA. SN: phage-enriched supernatant, Vol: sample volumes. **d-f)** Impact of other chemical and physical factors involved in purification on ssDNA recovery and quality. Agit: agitation, Inc: incubation. **g)** Pearson correlation between initial sample availability and final recovery and DNA quality. **h)** Prediction of phage genome recovery from the initial sample amount by linear regression. Data are presented as mean ± SD of three independent biological replicates (n=3). Statistical significance was assessed by one-way ANOVA followed by Tukey’s post hoc test for comparisons among three or more groups, and by unpaired t test with Welch’s correction for comparisons between two groups. Pearson correlation between SN volume and total DNA (r=0.9983, 95% CI 0.9728 to 0.9999, p<0.0001) or A_260/230_ ratio (r = 0.9204, 95% CI 0.2029 to 0.9948, p=0.0266) was calculated from the five mean values (**g**). Linear regression of total DNA yield and _A260/230_ ratio against SN volume (ml) gave Y = 14.63X + 0.4918 (R² = 0.9901) and Y = 0.9275X + 0.9804 (R² = 0.7985), respectively; both slopes were significantly non-zero (F(1, 13) = 1297 and 51.52, p<0.0001), n=5 (h). ns: not significant, *: p<0.05, **: p<0.01, ***: p<0.001.

### Downstream analysis of the Pf4 ssDNA genome provides complementary information to its intracellular dsDNA RF

Genome purification from filamentous phages should be useful for validating their identity by restriction patterns and PCR, and for straightforward sequencing. Hence, Pf4 ssDNA purified by all methods was tested for its utility in all these downstream applications. Both DNeasy and chaotropic-based methods gave the same assembled 7083 bp ssDNA sequence, with >200x coverage and a histogram centered around 7 kb (**Figure 4a-c**). On the other hand, sequencing of the phage dsDNA RF showed an assembled 6968 bp genome, lacking 115 bp in the PA0724-gIII gene relative to the viral ssDNA (**Figure 4d**). A closer look at the NGS histograms reveals two intracellular RF forms and a broad ssDNA peak, indicating that at least two phage genomes, the 6968 and 7083 bp sequences, coexist and are produced by the Pf4 phage. This kind of Pf4 genetic instability has been well documented previously under different experimental conditions^37,74,75^. In our studies, the Pf4 variants were detected after one round of amplification, suggesting rapid phage evolution *in vitro*, although their stable coexistence in the long term remains unknown.

**Figure 4.**
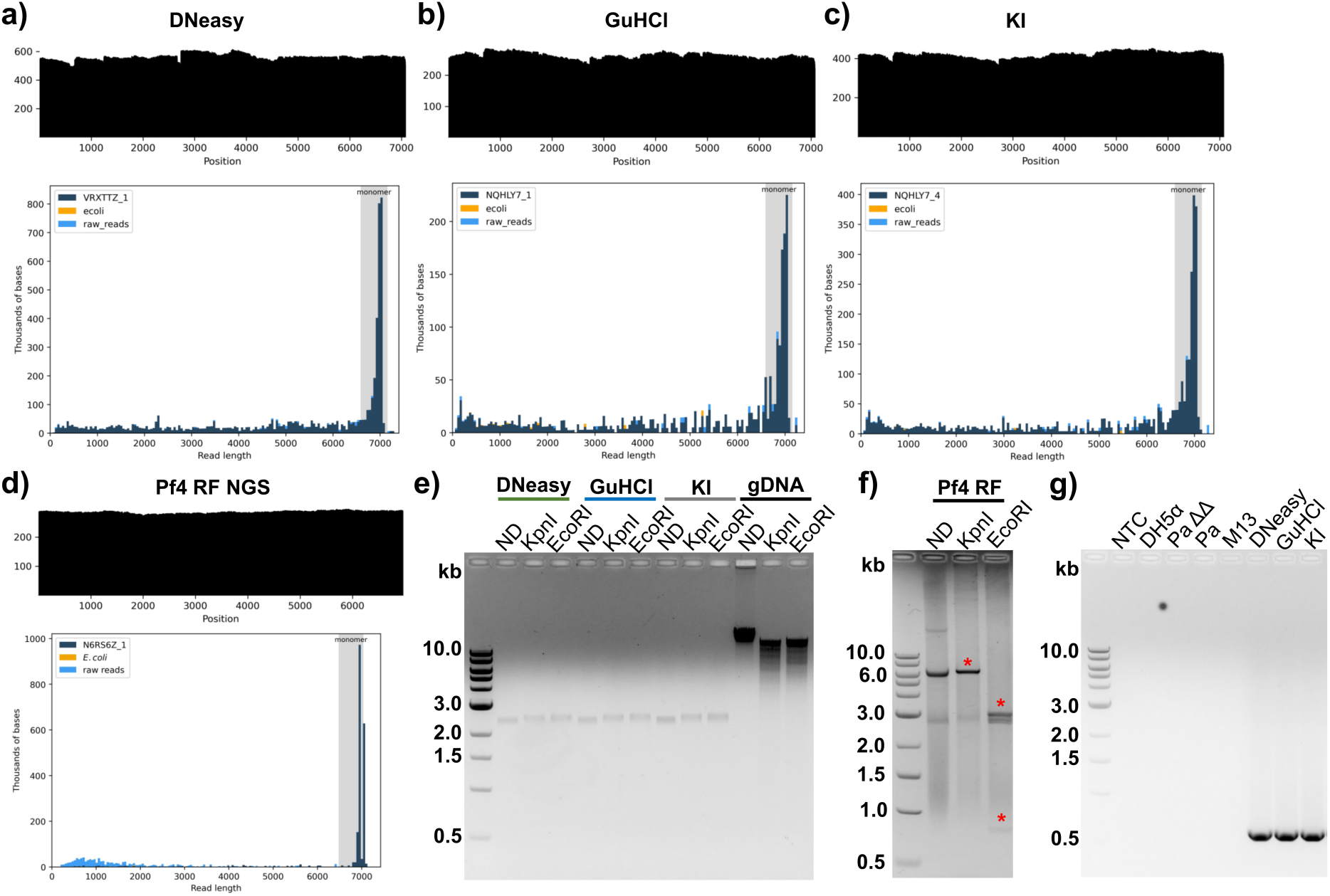
Downstream applications of viral ssDNA purified with the commercial DNeasy Blood & Tissue kit and chaotropic salt-based protocols. **a-c)** Next-generation sequencing (NGS) of purified ssDNA, showing coverage plots (top) and histograms (bottom) for each protocol. **d)** NGS of the Pf4 replicative form (RF) purified from infected cells with a plasmid miniprep kit, showing the coverage plot (top) and histogram (bottom). **e)** Restriction analysis of ssDNA purified with all protocols, visualized on a 1% agarose gel. 200 ng of total DNA was applied per lane alongside a 1 kb DNA ladder. gDNA: gDNA purified from the MPAO1 strain as a positive dsDNA control for digestion. ND: undigested control. **f)** Restriction analysis of purified Pf4 dsDNA RF with *KpnI* and *EcoRI*. 2 μg of total DNA was applied per lane on a 1% agarose gel. Red asterisks indicate the expected linear Pf4 7 kb genome (*KpnI*) and the 3117, 2832 and 806 bp bands (*EcoRI*). 213 bp band from EcoRI is not visible/marked. **g)** PCR amplification of the Pf4-specific gII-PA0717 region from purified Pf4 ssDNA, alongside a 1 kb DNA ladder and negative controls (NTC: non-template control, DH5α: cell lysate from *E. coli* DH5α strain, Pa ΔΔ: cell lysate from *Pseudomonas aeruginosa* MPAO1 ΔPf4 ΔPf6 strain, Pa: cell lysate from MPAO1 strain, M13: purified ssDNA from the *E. coli* phage M13KO7). kb: kilobases.

Restriction digestions of phage DNA help confirm viral identity and ssDNA conformation, and can also be used to eliminate contaminating host genomic DNA from samples. The restriction patterns for ssDNA (**Figure 4e**) and RF (**Figure 4f**) also showed characteristic features. While *KpnI* and *EcoRI* were able to digest the dsDNA RF, resulting in the expected linear genome and three ∼800-3000 bp bands, respectively, these enzymes failed with the purified ssDNA for all protocols. The fourth 213 bp band from *EcoRI* digestion of Pf4 RF was not visible in the gel. The upward shift observed in the agarose gel for enzyme-treated genomes in comparison to the non-digested ssDNA (ND) is consistent with these enzymes binding to their recognition sites but failing to cut. Given that the control dsDNAs (phage RF and MPAO1 gDNA) were successfully digested, and that the REBASE database (Accessed on October 5_th_, 2026) reports 60% and 40% of residual activities for *KpnI* and *EcoRI* on linear 42 bp and 46 bp oligonucleotides, respectively, their failure on Pf4 ssDNA is likely related to substrate conformation.

Enzyme activity on the REBASE oligonucleotides can be explained only by double-stranded structures containing the restriction sites and formed by dimerization. Indeed, the lowest-free-energy hybridization structures calculated by the DINAMelt server showed that oligonucleotides tested on *KpnI* and *EcoRI* enzymes have a T_m_ 6-11°C above the 37°C digestion temperature (**Supplementary Table IV**). Therefore, most of these oligos are in a transient conformation in which the recognition sites are located within the paired regions and accessible to enzymes, consistent with residual activities reported on ssDNA (**Supplementary Figure 4**). The same conclusions have been drawn for *HhaI* and *CfoI* enzymes, where oligonucleotides having their recognition sites within high stability dimers with completely paired forms and adjacent bases were efficiently cleaved_76_.

Thermodynamic calculations for local structures around *KpnI* and *EcoRI* sites in the Pf4 genome did not show any of their recognition sites to be paired according to our predefined criteria (**Supplementary Figure 5**). A closer inspection of these sites reveals that they are partially paired and adjacent to stable stems, the former explaining the lack of activity on this substrate and the latter their binding. The existence of local secondary structures in filamentous phage genomes has been demonstrated before for M13, allowing ssDNA cleavage depending on the enzyme _77_. However, their stability correlates with the overall cleavage rate and influences the rate of enzymatic reaction by favoring some DNA-enzyme complexes over others_76_.

Successful PCR amplification of the internal Pf4 gII-PA0717 region confirmed that no PCR-interfering contaminants are present in the final ssDNA from any of the DNeasy or chaotropic-based protocols, validating their use for this application (**Figure 4g**). The absence of any amplified band in the *E. coli* filamentous phage M13 and in the *E. coli* K12 and Pa strains free of this Pf4 variant confirmed the specificity of this PCR.

### Higher performance of chaotropic-based protocols extends to M13KO7, a second filamentous phage

These chaotropic-based protocols were developed for the filamentous phage Pf4 and optimized for the purification of phage genomic ssDNA. To test whether they extend to other filamentous phages, we applied these protocols to a second filamentous phage from *E. coli*, M13KO7, and compared them with the DNeasy kit. M13KO7 phage was propagated to 1.5 ± 0.2 x 10^11^ PFU/ml, a titer in the same 10^10^-10^12^ PFU/ml range commonly reported for M13 and related filamentous phages from *E. coli* ^8,24,30^. Nanodrop again overestimated ssDNA relative to fluorescence for M13KO7 for DNeasy (p=0.0131) and GuHCl (p=0.0327), but not for KI protocol (p=0.0534). Both DNA measurement techniques showed that chaotropic-based protocols recovered 2.2-2.6 times more DNA than the DNeasy kit (**Figure 5a**, Nanodrop: DNeasy vs GuHCl and KI p=0.0001 and p=0.0002, respectively; Fluorescence: DNeasy vs GuHCl and KI p=0.0017 and p=0.0034, respectively).

**Figure 5.**
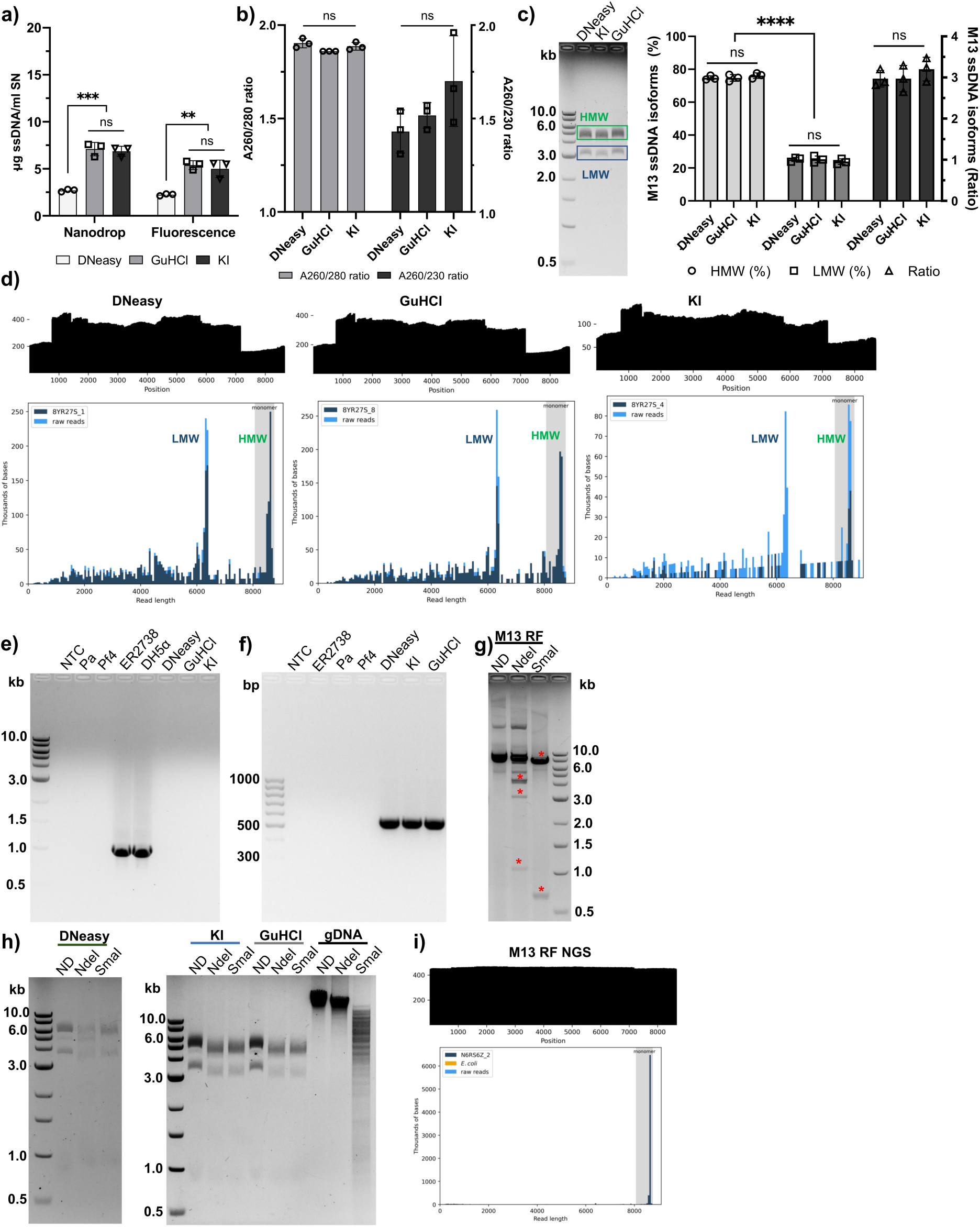
Application of the commercial DNeasy Blood & Tissue kit and chaotropic-based protocols to ssDNA purification from the *E. coli* filamentous phage M13KO7. **a-b)** ssDNA recovery and quality control (A_260/280_ and A_260/230_ ratios) of purified ssDNA. **c)** Evaluation of ssDNA integrity and purity of all purification protocols on a 1% agarose gel. 500 ng of total DNA was applied per lane alongside a 1 kb DNA ladder. The percentage represented by each band/isoform (HMW: high molecular weight; LMW: low molecular weight) and the ratio between them were calculated in ImageJ software. **d)** NGS of purified ssDNA, showing coverage plots (top) and histograms (bottom) for each protocol. **e)** Detection of contaminating gDNA from the ER2738 host strain and the related *E. coli* K-12 DH5α in M13 ssDNA purified with the DNeasy, GuHCl and KI protocols, by PCR targeting the K-12 rfb locus. **f)** PCR amplification specific for the M13KO7 gIII gene, alongside a 1 kb DNA ladder and negative controls (NTC: non-template control, ER2738: cell lysate from the *E. coli* host ER2738, Pa: cell lysate from MPAO1 strain, Pf4: purified ssDNA from Pf4 phage). kb: kilobases. **g)** Restriction pattern of purified M13KO7 dsDNA replicative form (RF) with *NdeI* and *SmaI*. 2 μg of total DNA was applied per lane on a 1% agarose gel. Red asterisks indicate the expected 4462, 3127 and 1080 bp bands (*NdeI*) and 7980 and 689 bp bands (*SmaI*). **h)** Restriction patterns of purified M13KO7 ssDNA with *NdeI* and *SmaI* enzymes. 500 ng of total DNA was applied per lane on a 1% agarose gel. gDNA from the MPAO1 strain was used as a dsDNA control for digestion. ND: undigested. **i)** NGS of the M13KO7 RF purified from infected cells with a plasmid miniprep kit, showing the coverage plot (top) and histogram (bottom). Data are presented as mean ± SD of three independent biological replicates (n=3). Statistical significance was assessed by one-way ANOVA followed by Tukey’s post hoc test for comparisons among three or more groups, and by unpaired t test with Welch’s correction for comparisons between two groups. ns: not significant, *: p<0.05, **: p<0.01, ***: p<0.001, ****: p<0.0001.

Regarding quality, all protocols resulted in high A_260/280_ and A_260/230_ ratios with no significant differences between them (**Figure 5b**, A_260/280_ p=0.0777, A_260/230_ p=0.1889). Also, two ssDNA bands, with no evidence of DNA degradation in any case, were observed in an agarose gel (**Figure 5c**). The relative abundance of the purified M13 ssDNA bands, labeled as low-molecular-weight (LMW) and HMW, estimated by ImageJ as HMW:LMW ratios, was around 3:1 across all protocols (**Figure 5c**, HMW vs LMW p<0.0001 for all three protocols). NGS of these products confirmed the co-purification of both M13 forms in all cases, with sizes of 6406 and 8669 bp, >200X coverage and well-defined peaks in the histograms (**Figure 5d**). In parallel, the absence of contamination from the bacterial host, assessed by a PCR specific to *E. coli* K12 strains, was confirmed for all purification strategies (**Figure 5e**). The phage genomes could also be used successfully as a template to amplify the M13 gIII gene (**Figure 5f**).

The generation of characteristic restriction patterns for the phage DNA was also applied to the M13KO7 phage using the *NdeI* and *SmaI* enzymes. The ability of these enzymes to cleave at their specific sites was initially tested on miniprep-purified dsDNA M13KO7 RF (**Figure 5g**), showing the expected 4462, 3127 and 1080 bp linear bands for *NdeI* digestion, and 7980 and 689 bp for *SmaI*. Next, their residual activities on ssDNA, reported in the REBASE database as 70% and 10% for *NdeI* and *SmaI*, respectively, were correlated with the best thermodynamic structures generated by the DINAMelt server (**Supplementary Figure 6**). For the *NdeI* oligonucleotide, the restriction site was not located in any double-stranded transient structure, which predicts no activity on this substrate. However, theoretical competing structures with a similar stability have been calculated for M13 ssDNA, explaining the 70% activity reported for this enzyme. For *SmaI*, on the other hand, the low activity on oligonucleotides can be ascribed to the two imperfectly paired restriction sites theoretically predicted by thermodynamic calculations (**Supplementary Figure 6b**). Although both enzymes were active on dsDNA MPAO1 gDNA, no cleavage was observed for any purified phage ssDNA in an agarose gel (**Figure 5h**). Predicted structures around these restriction sites showed their location in non-paired or partially paired sequences (**Supplementary Figure 7**), in agreement with their lack of activity on this ssDNA substrate and the requirement of stable double-stranded structures for efficient cleavage^76^.

### Analyzing both RF and ssDNA reveals selective deletion revertants of M13KO7 at the ssDNA level

M13 phage has been extensively used as a phage vector for DNA cloning and peptide display_5,8_, and as a helper phage to produce encapsulated phagemids and for peptide/protein display purposes as well ^78,79^. The M13KO7 helper phage was made by inserting a kanamycin resistance gene and a low-copy p15A origin of replication at the *AvaI* site in the M13 IG ^31^. As this insertion makes the phage less efficient during replication, phagemids are preferentially packaged into virions at approximate ratios of 1:10 – 1:100 (M13KO7:phagemid), making M13KO7 a powerful tool for phagemid production^29,31,80^.

Unexpectedly, after purification, agarose gel visualization and sequencing of the M13KO7 ssDNA revealed the production of two different phage populations, one having the original 8669 bp sequence reported for this phage vector and a second, shorter 6406 bp form differing only by a 2263 bp deletion (**Figure 5c, d, h**). This deletion includes the whole kanR-p15ori insert and restores the wild-type M13 origin. To elucidate whether this deletion originated at the level of ssDNA replication/processing or in the phage RF, the RF was miniprep-purified and sequenced, showing an 8669 bp sequence with a single peak in the histogram and a >400x coverage (**Figure 5i**).

The presence of only one RF population can also be inferred from its digestion with *NdeI* and *SmaI* (**Figure 5g**). Both enzymes efficiently digested the RF, yielding the bands expected for M13KO7, and were fully active on the control Pa gDNA (**Figure 5h**). Therefore, unlike the ssDNA, where secondary structures can hinder cleavage (**Figure 4e**), the dsDNA RF shows no evidence of inhibited digestion. Although *NdeI* is known for partial digestions, only the bands corresponding to M13KO7 and undigested dsDNA were observed, and the 2200 bp band predicted for the shorter phage was absent. *SmaI* gave the same result: only the expected M13KO7 bands were present. Since the SmaI site is not included in the shorter phage RF, an undigested ∼6.4 kb band would be expected if the 6406 bp RF coexisted with the full-length vector, and no such band was detected. The absence of the bands expected from the shorter phage is thus consistent with the lack of the shorter M13KO7 form at the RF level.

### Chaotropic-based purification recovers 60-100% of the total ssDNA present in PEG-precipitated phage samples

The same or better phage-dependent performance of the chaotropic-based protocols in comparison with commercial gDNA extraction alternatives such as Monarch and DNeasy kits was shown in the previous sections. However, how much of the total input phage ssDNA these methods could recover remained unknown. This metric helps to determine protocol efficiency for different filamentous phages and, since input ssDNA correlates significantly with recovery (**Figure 3g-h**), it could be used to anticipate expected phage DNA concentrations and recovery case by case. Hence, we proceeded to quantify total phage ssDNA in the SN before purification and in the purified sample using the Qubit ssDNA Assay kit (**Figure 6a-b**). DNeasy kit and chaotropic-based protocols recovered the same ∼11 μg ssDNA/ml SN for Pf4 phage (p=0.1507), which corresponded to ∼100% of the ssDNA measured in the PEG-precipitated samples. M13KO7, on the other hand, had a significantly higher recovery when employing chaotropic-based protocols than the DNeasy kit (DNeasy vs GuHCl p=0.0001, DNeasy vs KI p=0.0002), as seen before. However, from an input of ∼8 μg/ml SN, the purified ssDNA dropped to ∼5 μg/ml SN for the chaotropic-based protocols (GuHCl p=0.0066, KI p=0.0026) and to ∼2.3 μg/ml SN for the DNeasy kit (p<0.0001), representing about 60% and 28% of the total input, respectively.

**Figure 6.**
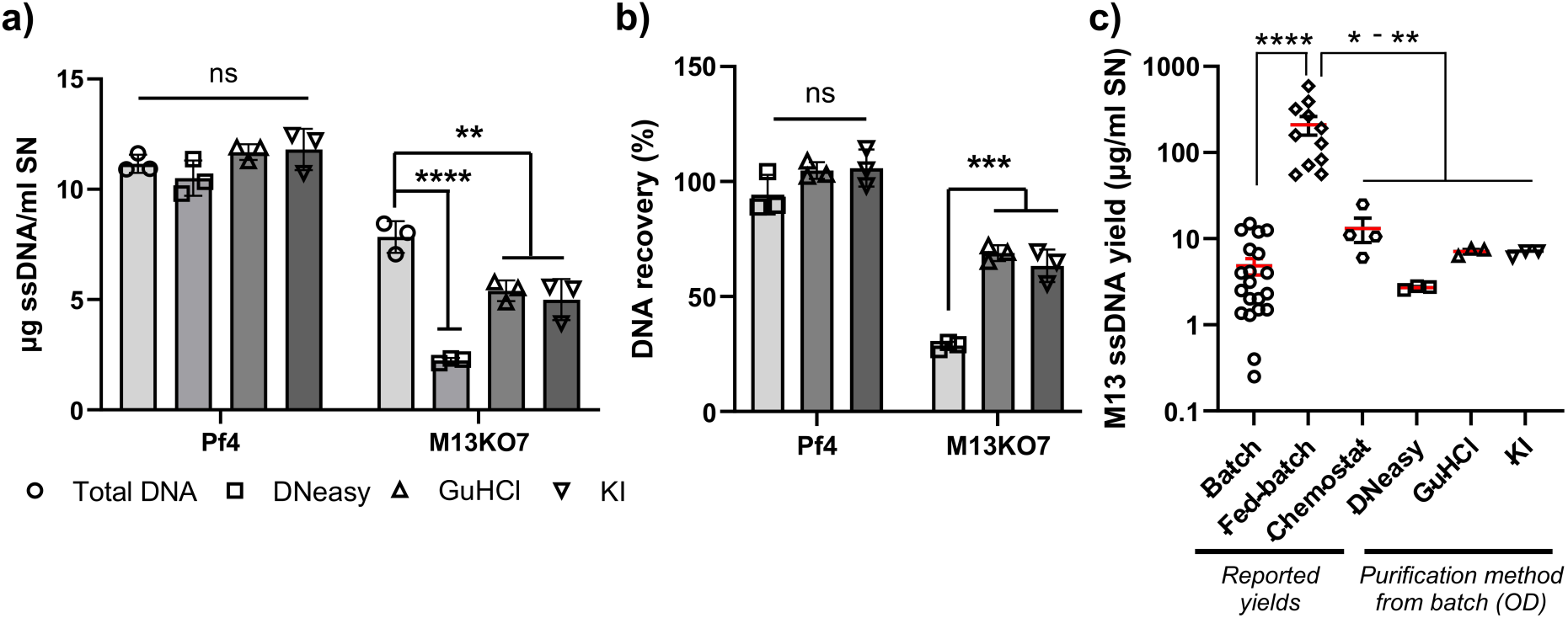
ssDNA recovery with the commercial DNeasy Blood & Tissue kit and chaotropic-based protocols. **a-b)** Comparison of ssDNA recovery from the filamentous phages Pf4 and M13KO7 between the three protocols and the total ssDNA available in the sample, measured by fluorescence. **c)** Comparison of M13KO7 ssDNA yield from the three characterized protocols with values reported in the literature for M13 phage in batch, fed-batch or chemostat bioprocesses. Data are presented as mean ± SD of three independent biological replicates (n=3). Statistical significance was assessed by one-way ANOVA followed by Tukey’s post hoc test for comparisons among three or more groups, and by unpaired t test with Welch’s correction for comparisons between two groups. ns: not significant, *: p<0.05, **: p<0.01, ***: p<0.001, ****: p<0.0001.

The efficiencies of the chaotropic-based and DNeasy methods were compared with literature reports in which ssDNA recovery per volume of SN was available or could be calculated from the data (**Supplementary Table V**). As ssDNA recoveries have not been reported before for Pf4, we decided to use wild-type M13 and all derivative vectors as a reference. Published studies encompassed phage propagation in batch, fed-batch, and chemostat (continuous) bioprocesses (**Figure 6c**), and a very broad variety of techniques, such as direct quantification in crude phage lysates or after ethanol precipitation, classical phenol/chloroform extraction, capture by magnetic beads or glass fiber plates, or plasmid miniprep and midiprep kits. Both chaotropic-based and DNeasy protocols were in the same range as batch and chemostat (p>0.9999), as expected from our cultures in batch mode, and significantly lower than fed-batch (DNeasy p=0.0089, GuHCl p=0.0110, KI p=0.0109). The very high bacterial biomass reached in fed-batch processes (20-94 g/L) translated into a viral concentration of 10_13_ – 10_15_ PFU/ml in the SN, consistent with higher ssDNA yields^52,81–84^.

### ssDNA recovery on silica is phage-dependent and not limited by electrostatic repulsion

Following up on the lower recovery of M13KO7, we decided to enhance the molecular interactions between phage ssDNA and silica in order to increase DNA yield. Interactions governing DNA binding to silica can be classified into repulsive (electrostatic interactions) and attractive (hydrophobic interactions and hydrogen bonding) forces^65^, where the repulsive forces could be weakened by decreasing the pH at which DNA binding occurs. At lower pH, the surface charge density of silica becomes less negative ^85^, which brings both interacting partners closer and favors attractive forces. A higher recovery at lower pH has been widely reported for dsDNA from very diverse sources, such as λ (48.5 kb), T7 (39 kb) and PSP3 (30.6 kb) phages^70,86,87^, pUC18 plasmid and salmon DNA ^71,88^, and ancient DNA from bones and teeth ^69^. We therefore tested whether lowering the pH of DNA binding would increase ssDNA recovery.

Pf4 and M13KO7 phage genomes were purified by the KI-based protocol with two different dissolving buffers: PBS (pH 7.4) and acetate (pH 5.0). In the PBS protocol, the pH decreased to ∼7.0 after KI addition, returned to ∼7.5 for DNA binding and was 8.5 for elution. In the acetate protocol, the pH was 5.0 during phage solubilization and disaggregation, rose to ∼6.0 for DNA binding and was 8.5 for elution (**Figure 7a**). Fluorescence quantification of Pf4 ssDNA showed no differences in recovery (**Figure 7b**, p=0.1526) or DNA quality (**Figure 7c**, A_260/280_ p=0.1296, A_260/230_ p=0.1539) for any buffer/pH, while a lower pH led to a significantly lower yield for M13KO7 (**Figure 7d**, Nanodrop p=0.0058, Fluorescence p=0.0026) without affecting quality (**Figure 7e**, A_260/280_ p=0.5735, A_260/230_ p=0.1430).

**Figure 7.**
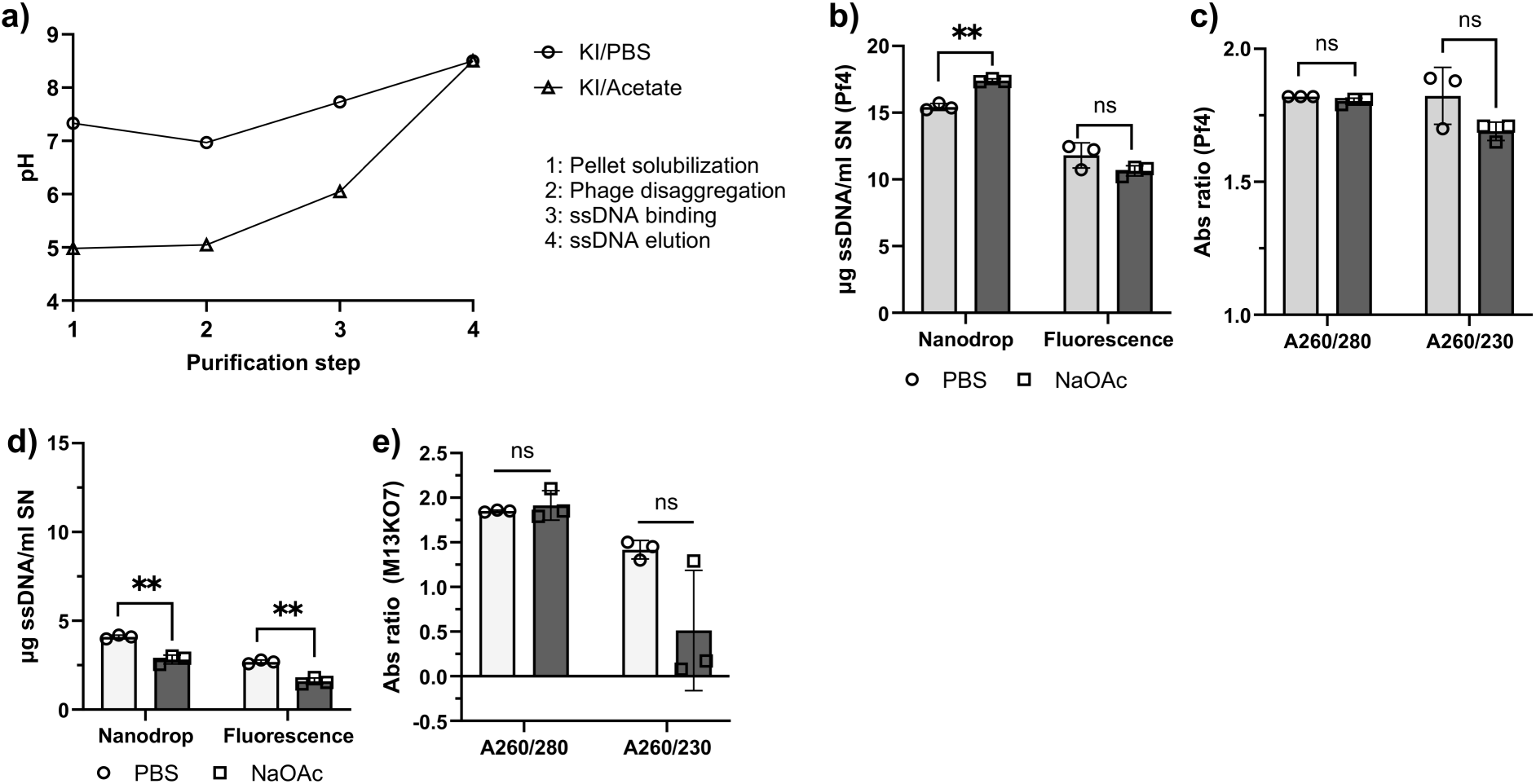
Evaluation of ssDNA-silica interactions at different pH values for Pf4 and M13KO7 phages. **a)** Measurement of the pH at each step of ssDNA purification with the chaotropic salt KI when the sample was solubilized in PBS or acetate buffer. **b-c)** Pf4 ssDNA quantification (Nanodrop and fluorescence) and quality (A_260/280_ and A_260/230_ ratios) when the phage pellet was solubilized in PBS or acetate buffer prior to KI-based purification. **d-e)** M13KO7 ssDNA quantification (Nanodrop and fluorescence) and quality (A_260/280_ and A_260/230_ ratios) when the phage pellet was solubilized in PBS or acetate buffer prior to KI-based purification. Data are presented as mean ± SD of three independent biological replicates (n=3). Statistical significance was assessed by unpaired t test with Welch’s correction for comparisons between two groups. ns: not significant, *: p<0.05, **: p<0.01.

Lowering the pH at the DNA-silica binding step from ∼7.5 to ∼6.0 therefore did not improve ssDNA recovery for Pf4 and reduced it for M13KO7. This suggests that dehydration by chaotropic salt and ethanol is sufficient to overcome electrostatic repulsion in the pH range tested, so repulsion does not appear to limit ssDNA binding to silica under these conditions. The phage-specific decrease for M13KO7 indicates that the response to pH depends on the phage.

## Discussion

The development and optimization of purification protocols for genomic DNA has focused mainly on the double-stranded members of these viruses ^39–49^. Consequently, we have optimized and thoroughly characterized two easy-to-implement and fully customizable GuHCl/KI-silica columns-based protocols to purify and analyze genomic ssDNA from filamentous phages. As commercial kits based on silica columns are the preferred method for phage purification, we compared these protocols side by side with two commercial alternatives, the Monarch Spin gDNA Extraction and DNeasy Blood & Tissue kits. The filamentous phages Pf4 and M13 were chosen to represent this group because of their biotechnological and clinical importance.

Among the commercial alternatives, the DNeasy kit consistently gave about 6.6 times higher recovery than the Monarch kit, which is in agreement with their different primary purposes; while Monarch was developed for high-molecular-weight genomic DNA, DNeasy is intended to recover most DNA from any source in samples. The chaotropic-based protocols recovered the same or higher phage genome yields than the DNeasy kit, while maintaining the same quality and suitability for many downstream applications such as NGS, PCR and restriction analyses. The experimental restriction patterns were dependent on the formation and stability of secondary structures in the purified ssDNA, and as these depend on phage sequence are phage specific and could help to corroborate predicted local ssDNA structures and contribute to validating phage identity, as other ssDNA substrates could be cleaved depending on their local folding.

These protocols recover 60-100% of total ssDNA and, in the two phages tested, varied less between phages than the DNeasy kit. We hypothesize that the phage-dependent differences in recovery could arise from different GC contents (62% Pf4 vs 43% M13KO7) and sequence-dependent secondary structures in the ssDNA, which modulate its interaction with the silica matrix. Compared with previously reported ssDNA purification methods, recovery from chaotropic-based protocols was comparable, with the added advantages of using standard silica columns and of being customizable. They are also independent of silica column suppliers and fully customizable, unlike commercial alternatives where many solutions are proprietary and cannot be adjusted for specific phages. The GuHCl/KI salts selected for the present protocols are extremely soluble, easy to prepare as concentrated solutions, less viscous, and cheaper than stronger chaotropic alternatives. GuSCN, a very common and stronger chaotropic salt in commercial kits, is classified as harmful to aquatic life with long-lasting effects ^89^, and when in contact with strong oxidizers can generate highly toxic byproducts, including cyanide-related species under some conditions_90_.

When chaotropic-based procedures are implemented for the study of the biology of filamentous phages through ssDNA purification, in combination with plasmid miniprep kits for the purification of the phage RF, they give complementary and very useful information. While ssDNA from Pf4 phage confirmed the mutations and deletions observed in the double-stranded RF, M13 ssDNA sequencing and analyses suggested a possible mechanism by which shorter, faster-propagating variants arise. NGS of both RF and ssDNA and RF digestion indicated that under selective pressure the RF keeps the genetic elements necessary to survive while the ssDNA, packaged into virions and playing no role in bacterial fitness, is subject to selection for phage fitness and genome reduction. Whether this variation affects the performance of helper phages such as M13KO7 (for example, through helper-phage carry-over in phagemid preparations) remains to be tested, but it may be relevant when engineering other filamentous phages. The local folding and flexible conformations of ssDNA can also be used to confirm these structures experimentally when predicted *in silico* and contribute to the identification and validation of several phages. We therefore recommend the implementation of RF and genomic ssDNA purification, sequencing and analyses in common workflows for the study of filamentous phages.

The genetic drift observed for Pf4 phage in both the RF and ssDNA correlates with previous reports about its genetic instability. Pf4 phages produced by the MPAO1 strain experimentally evolved for 1000 generations *in vitro* showed a hyperactive phenotype, with mutations in the repressor region, non-synonymous nucleotide polymorphisms in structural and functional genes, and deletions upstream of pf4r, the toxin-antitoxin module and the integrase gene ^74^. In a flow-biofilm model, 8-day-old MPAO1 cultures released ∼10^11^ PFU/ml of super-infective phages, Pf4 variants capable of infecting lysogenized and non-lysogenized Pa with the same efficiency. These phages mostly carried a 3.4 kb deletion upstream of the excisionase gene in a process dependent on the prophage phrD and pfrT genes ^37^. Besides the known genetic drift of Pf4 phages, the stable coexistence of different Pf4 forms inside the same cell has also been documented in PAO1 cells infected with recombinant Pf4 phages carrying carbenicillin or tetracycline resistance genes inserted between the pfiT gene and the attR site ^75^. Phage production was, in this case, induced by the expression of Pf4 excisionase under an arabinose promoter, and daughter cells exhibited a double-resistant phenotype in the absence of antibiotic selection.

Differently and to the best of our knowledge, the only antecedents showing similar genome mutations and reductions in engineered M13 phages are the compensatory mechanisms developed by the M13KE vector to restore wild-type propagation rates^27,91,92^. M13KE is a cloning/display vector carrying a polylinker within the lacZα gene in the IG with the same deleterious consequences for replication as M13KO7. This phage is an M13mp19 derivative in which the N-terminal end of the gIII gene has been engineered for peptide display ^93^. The compensatory mechanisms include more than a dozen mutations in the 5’UTR of the gII gene, a silent mutation in the gene IV coding sequence, and the ejection of most of the lacZα insert. Unlike M13KO7, M13KE-based phage Ph.D. libraries are generally stable, and, although not explicitly stated for all mutations, the genetic differences in the ssDNA from fast-propagating clones in comparison with wild-type M13KO7 were re-confirmed in the RF DNA^27^.

These findings are consistent with a model of M13KO7 propagation in which the RF is maintained unaltered within the cell, whereas shorter, faster-propagating variants arise alongside full-length genomes at some stage of ssDNA synthesis, processing, export or packaging (**Figure 8**). We hypothesize that selection for phage fitness acts mainly on the virion ssDNA. Because the M13 replication origin is compromised, variants with improved replication, packaging or secretion can emerge during phage amplification, consistent with deletion of the p15A ori-kanR insert (**Figure 8a**). These variants represent about 25% of the purified ssDNA by gel band intensity, indicating that the full-length genome still predominates (**Figure 8b**). This is expected, since neither phage population offers a selective advantage in a culture where every cell already carries the full-length RF, maintained by kanamycin selection. Reinfection would not confer any new fitness benefit, and any RF derived from the shorter variants would be lost, as it would lack the kanR cassette and could not compete with the full-length RF under antibiotic selection (**Figure 8c**).

**Figure 8.**
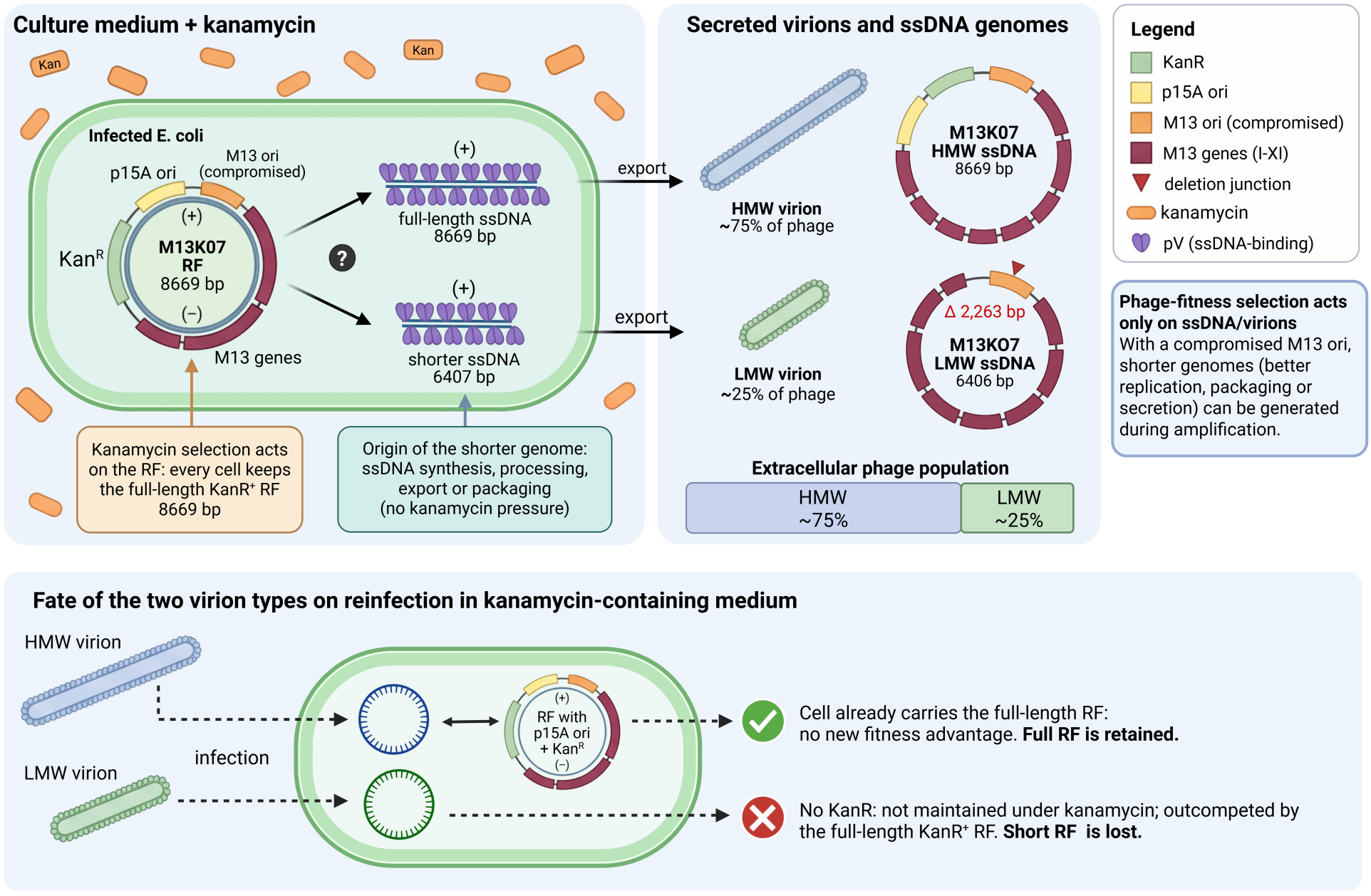
Proposed mechanism for the generation of M13KO7 deletion-revertant phages. **a)** Intracellular events. The double-stranded replicative form (RF; 8669 bp) of M13KO7 carries the p15A origin, the kanamycin resistance gene (KanR), the compromised M13 origin and the M13 genes (I-XI). The full RF is maintained in every cell by kanamycin selection. Plus-strand ssDNA is synthesized from the RF and coated by pV, yielding full-length (8669 bp) and shorter (6406 bp) genomes. The question mark indicates that the step at which the shorter genome arises is unresolved (ssDNA synthesis, processing, export or packaging); no kanamycin pressure acts at this stage. **b)** Extracellular products. Both genome types are exported and packaged into virions. The high-molecular-weight (HMW) virions contain the full-length 8669 bp ssDNA and the low-molecular-weight (LMW) virions contain the 6406 bp ssDNA, in which the p15A ori-KanR insert is deleted (Δ2263 bp; red triangle marks the deletion junction). Virion lengths and genome maps are drawn in proportion to genome size. The stacked bar shows the approximate proportion of each virion type in the extracellular phage population, estimated from the relative band intensities of the purified ssDNA on agarose gels (ImageJ; Fig. 5c). Phage-fitness selection is proposed to act mainly on the ssDNA/virions; because the M13 origin is compromised, shorter genomes with improved replication, packaging or secretion can be generated during amplification. (**c**) Fate of the two virion types upon reinfection in kanamycin-containing medium. Cells already carry the full-length RF, so reinfection by HMW virions offers no new fitness advantage and the full-length RF is retained. LMW virions give rise to an RF lacking the p15A ori and KanR, which cannot be maintained under kanamycin and is outcompeted by the full-length KanR⁺ RF, so the shorter RF is lost. Kan, kanamycin; ssDNA, single-stranded DNA. Created in BioRender. Perez Rodriguez, S. (2027) https://BioRender.com/ldmn42o.

The key role of chaotropic salts and ethanol identified in silica-based ssDNA purification can be explained by a water-exclusion model^65,88^. ssDNA binding to silica is regulated by three different forces: hydrophobic interactions and hydrogen bonding, which favor DNA binding to silica silanols and the hydrophobic surface, and electrostatic interactions, which lead to repulsion between the negative charges of DNA and silica. In the absence of chaotropic salts and ethanol, DNA and silica are surrounded by a hydration shell that impedes short-range interactions (**Figure 9a**). Chaotropic salts promote DNA binding to silica by perturbing the hydrogen-bonding network of water. These salts disrupt the ordered water structure and strip the hydration shell from the DNA phosphate backbone and silica surface, facilitating DNA-silica interactions ^71,72^. Under these dehydrating, high-ionic-strength conditions, DNA can interact with the silica surface through hydrophobic forces, hydrogen bonding and cation-mediated bridging (**Figure 9b**)^65,86,87^. Adding ethanol to samples increases this dehydration and favors DNA binding ^88^. Molecular dynamics simulations predict that hydrophobic interactions and hydrogen bonding contribute equally to binding ^65^, and experimental data strongly support sample dehydration as the mechanism guiding DNA binding to silica surfaces^70–72,86,87^.

**Figure 9.**
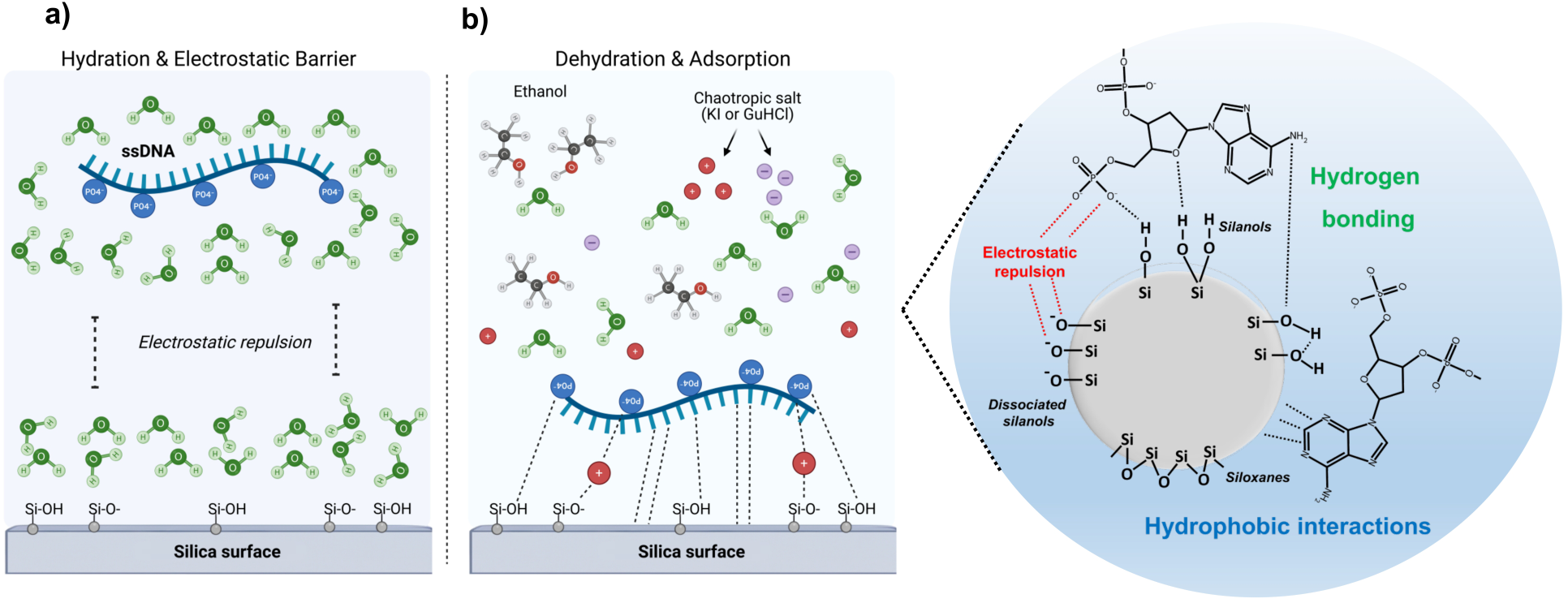
Proposed water-exclusion model for ssDNA-silica binding. **a)** DNA-silica interactions are initially limited to weaker long-range interactions because of the water shell solvating both molecules. Electrostatic repulsion between DNA phosphate groups and dissociated silanols on the silica surface further reduces DNA binding. **b)** Addition of ethanol and chaotropic salts disrupts the water shell and promotes short-range DNA-silica interactions by bringing the two closer. The high ionic concentration shields negative charges and decreases repulsive forces. **c)** Detailed diagram of the chemical groups available at the surface of a silica particle and their interactions with ssDNA (a nucleotide is represented for simplicity). Interactions were grouped into those promoting binding (hydrogen bonding and hydrophobic interactions) or repulsion (electrostatic). Created in BioRender. Perez Rodriguez, S. (2027) https://BioRender.com/ldmn42o.

Regarding electrostatic repulsion, our ssDNA purifications using chaotropic salts under neutral and acidic pH suggest that these interactions do not limit ssDNA binding to silica, supporting hydrophobic interactions and hydrogen bonding as the main forces in DNA-silica interactions. Unlike dsDNA ^70,71,86–88^, we found that a lower pH does not improve recovery of ssDNA and could, in some phage-specific cases, be detrimental. As the ssDNA from filamentous phages is more flexible than dsDNA and can adopt different conformations and local secondary structures, its interaction at different pH values, and therefore its binding behavior, is difficult to predict. DNA structure has been shown previously to affect recovery for linear or circular supercoiled plasmids ^88^ and phage dsDNA with different non-paired end lengths ^86^. In conclusion, the pH effect seems to be related to phage sequence and the characteristic local structures formed.

The water-exclusion model could also be used to improve ssDNA recovery in difficult cases such as M13 phage. This model predicts that increasing sample dehydration should lead to higher binding and final recovery. Experimental evidence indicates that a higher concentration of the chaotropic salt will increase dehydration and favor DNA binding to silica^70–72^. Changing to a stronger chaotropic salt such as GuSCN may also improve recovery. However, the strength of DNA binding to silica should be regulated, as under experimental conditions in which DNA is tightly bound to this surface, elution is not efficient and recovery is negatively affected^70,71,88^. The application of these optimized chaotropic-based protocols to other filamentous phages will help to establish their broader applicability and advance the study of this important group of phages.

This study has limitations. Only two filamentous phages (Pf4 and M13KO7), with genomes of about 7.1 and 8.7 kb, were tested, so the recovery values may not hold for phages with other genome sizes, GC contents or secondary structures, and only two commercial kits were used for comparison. Recovery was calculated relative to the ssDNA measured by fluorometry in PEG-precipitated pellets; it therefore excludes losses during precipitation and depends on the accuracy of the Qubit ssDNA assay. The proposed binding mechanism is inferred from recovery data and could be enriched by direct measurements of DNA in all the fractions across the purification to differentiate between the efficiency of DNA binding and elution.

## Conclusions

The chaotropic-based protocols developed here recovered ssDNA with equal or higher yield than the Monarch Spin gDNA Extraction Kit and the DNeasy Blood & Tissue Kit, while producing DNA of high quality suitable for NGS, PCR and restriction digestion. They achieved 60% and 100% recovery for *E. coli* phage M13 and *Pseudomonas aeruginosa* phage Pf4, respectively, and, together with the dsDNA replicative form (RF), enabled the analysis of both forms across the phage life cycle. The results are consistent with a water-exclusion model of binding, in which hydrophobic and hydrogen-bonding interactions with silica overcome electrostatic repulsion. As ssDNA recovery is phage-specific, optimization can be done for lower-yield phages in future work, if necessary, which the fully customizable nature of these protocols facilitates. Application to additional filamentous phages should further establish their general utility and support the characterization and engineering of new phages.

## Acknowledgements

SPR was supported by the Stanford Maternal and Child Health Research Institute through a postdoctoral fellowship, and by the Cystic Fibrosis Research Institute (CFRI) through an Elizabeth Nash Memorial Fellowship. AK acknowledges National Institutes of Health grant K08 AI190313. P.L.B. discloses support from National Institutes of Health grant R01 HL148184-01; R01 AI12492093; R01 DC019965, K24 AI166718, P01AI1960471, and R01 EB038154 and support from the CFF and the CFRI. The contents are those of the authors and do not necessarily represent the views of the funding agencies.

## Data Availability Statement

The genome sequences of all *Pseudomonas aeruginosa* MPAO1 strains and the filamentous phages M13KO7 and Pf4 generated in this study have been deposited in the NCBI BioProject database under accession number PRJNA1538112 (https://www.ncbi.nlm.nih.gov/bioproject/PRJNA1538112). All other data supporting the findings of this study are available within the article and its supplementary materials.

## Author Contribution Statement

SPR contributed to the design of the research, developed the methods, collected and analyzed the data, and wrote the manuscript. AK prepared the Pf4 phage banks, purified the phages, and contributed to data analyses. CM contributed to the research design and data analyses, acquired funding, and co-supervised the project. PLB conceived and supervised the study, acquired funding, contributed to the research design and method development, analyzed data, and wrote the manuscript. All authors participated in manuscript corrections and read and approved the final manuscript.

