## Supplementary Material for "Purification of filamentous phage ssDNA genomes using Chaotropic salt/silica methods"

**Authors and Affiliations:**

Perez-Rodriguez, Saumel[1]; Khosravi, Arya[1]; Milla, Carlos[2]; Bollyky, Paul L[3].

[1] Division of Infectious Diseases and Geographic Medicine, Department of Medicine, Stanford University, Stanford, CA 94305, USA.

[2] Cystic Fibrosis Clinic, Department of Medicine, Lucile Packard Children's Hospital, Stanford University School of Medicine, Stanford, California, USA.

[3] Division of Infectious Diseases and Geographic Medicine, Department of Medicine, Stanford University, Stanford, CA 94305, USA. Electronic address:.

Saumel Perez-Rodriguez: https://orcid.org/0000-0003-3852-008X

Arya Khosravi: https://orcid.org/0000-0002-4958-3556

Carlos Milla: https://orcid.org/0000-0001-5515-3053

Paul L. Bollyky: https://orcid.org/0000-0003-2499-9448

### Supplementary Tables

**Supplementary Table I. Primer information and amplicon characteristics.**

| Primer name | Sequence (5' – 3') | Target | Amplicon (bp) | Source | Notes |
|---|---|---|---|---|---|
| Pf4F-Webb2004 | AGCAGCGCGATGAAGCAAT | Pf4 RF form | 839 | 1 | Does not detect the Pf4 prophage, only the RF |
| Pf4R-Webb2004 | TAGAGGCCATTTGTGACTGGA | | | | |
| GendelF-Rice2009 | ATCTTTCCCGGGCTTTACG | attL of Pf4 prophage | 806 | 2 | Amplifies the bacteria–prophage junction at the attL site |
| GendelR-Rice2009 | GCAATTGGCAAAGTGTTCGA | | | | |
| IntF4-F-Gavric2022 | TCGAATTCCGCTTCCATCAC | Internal Pf4 sequences | 1001 | 3 | Partially amplifies two Pf4-specific genes |
| IntF4-R-Gavric2022 | CCTGATGCTTGGTCAGGTACG | | | | |
| Pf4/PA14-F-Gavric2022 | CAATGGTCGTCACGCAGAAC | attR of Pf4 prophage | 943 | 3 | Amplifies the bacteria–prophage junction at the attR site |
| Pf4/PA14-R-Gavric2022 | CCGCTCAACCCGATCTAC | | | | |
| Pf4-Rev-Schmidt2023 | TCATTGGGAGGCGCTTTCAT | attR of Pf4 (if integrated) or 5' – 3' sites of Pf4 (bacterial genome when phage is | 2020 (Integrated) 2570 (phage lost) | 4 | Multiplex PCR. When the Pf4 prophage is present, only Rev-Fwd1 amplifies because the Rev-Fwd2 product is too large for the polymerase (14,966 bp). When Pf4 is lost, Fwd1 cannot bind and Rev-Fwd2 amplifies a |
| Pf4-Fwd1-Schmidt2023 | GGATATGGAGCGTGGTGGAG | | | | |
| Pf4-Fwd2-Schmidt2023 | AGTGGCGGTTATCGGATGAC | | | | |

| | | | | | |
|---|---|---|---|---|---|
| | | lost) | | | 2570 bp product. |
| Pf6-Rev-Schmidt2023 | AGGATCAATGACCAGTGGCG | attR of Pf6 (if integrated) or 5' – 3' sites of Pf6 (bacterial genome when phage is lost) | 1016 (Integrated) 1572 (phage lost) | [4] | Multiplex PCR. When the Pf6 prophage is present, only Rev-Fwd2 amplifies because the Rev-Fwd1 product is too large for the polymerase (13,642 bp). When Pf6 is lost, Fwd2 cannot bind and Rev-Fwd1 amplifies a 1572 bp product. |
| Pf6-Fwd1-Schmidt2023 | GTGATCCACGTGTCCAACAG | | | | |
| Pf6-Fwd2-Schmidt2023 | CCCAGTGCAGATGACTTGGT | | | | |
| Fw-Pf6-RF-Saumel2024 | CGGACTGAACCACTCTGACA | Pf6 RF form | 353 | This study | Does not detect the Pf6 prophage, only the RF |
| Rv-Pf6-RF-Saumel2024 | CGCTGTATCAGGTGAAGCAC | | | | |
| Pf4_mut_FW | CACTATCTCGCCAAGGGTTG | Pf4 phage | 592 | This study | Detects Pf4 ssDNA |
| Pf4_mut_RV | CGTGGAAGGTGCAAATCTCT | | | | |
| M13_pIII_FW | CCTCTCGACGGCACTTATCC | gIII gene, M13 phage | 531 | This study | Specific for the gIII gene of M13 phage; used to validate biological materials |
| M13_pIII_RV | CTGTAGCGCGTTTTCATCGG | | | | |
| K12IS-L | CGCGATGGAAGATGCTCTGTA | K-12 rfb locus | 969 | [5] | Identifies E. coli K-12 derivatives by a characteristic genetic rearrangement involving an IS5 insertion sequence |
| K12-R | ATCCTGCGCACCAATCAACAA | | | | |

**Supplementary Table II. PCR reaction conditions and amplification parameters.**

| Primers | Target | Reaction conditions | Cycling conditions |
|---|---|---|---|
| Pf4F-Webb2004<br>Pf4R-Webb2004 | Pf4 RF form | EconoTaq PLUS GREEN 1X Master Mix, 25 pmol primers. Template: 1 µl lysate or 50 ng phage ssDNA | Initial denaturation: 2 min 95°C.<br>Amplification (35 cycles):<br>Denaturation: 30 sec 95°C<br>Annealing: 30 sec 59°C<br>Extension: 1 min 30 sec 72°C<br>Final extension: 5 min 72°C |
| GendelF-Rice2009<br>GendelR-Rice2009 | attL of Pf4 prophage | EconoTaq PLUS GREEN 1X Master Mix, 50 pmol primers. Template: 1 µl lysate or 50 ng phage ssDNA | Initial denaturation: 2 min 95°C.<br>Amplification (35 cycles):<br>Denaturation: 30 sec 95°C<br>Annealing: 30 sec 64°C<br>Extension: 1 min 30 sec 72°C<br>Final extension: 10 min 72°C |
| IntF4-F-Gavric2022<br>IntF4-R-Gavric2022 | Internal Pf4 sequences | EconoTaq PLUS GREEN 1X Master Mix, 50 pmol primers. Template: 1 µl lysate or 30 ng phage ssDNA | Initial denaturation: 2 min 95°C.<br>Amplification (30 cycles):<br>Denaturation: 30 sec 95°C<br>Annealing: 30 sec 62°C<br>Extension: 1 min 30 sec 72°C<br>Final extension: 5 min 72°C |
| Pf4/PA14-F-Gavric2022<br>Pf4/PA14-R-Gavric2022 | attR of Pf4 prophage | EconoTaq PLUS GREEN 1X Master Mix, 50 pmol primers. Template: 1 µl lysate or 30 ng phage ssDNA | Initial denaturation: 2 min 95°C.<br>Amplification (30 cycles):<br>Denaturation: 30 sec 95°C<br>Annealing: 30 sec 59°C<br>Extension: 1 min 72°C<br>Final extension: 5 min 72°C |
| Pf4-Rev-Schmidt2023<br>Pf4-Fwd1-Schmidt2023<br>Pf4-Fwd2-Schmidt2023 | attR of Pf4 (if integrated) or 5' – 3' sites of Pf4 (bacterial genome when phage is lost) | EconoTaq PLUS GREEN 1X Master Mix, 25 pmol primers. Template: 1 µl lysate or 50 ng phage ssDNA | Initial denaturation: 2 min 95°C.<br>Amplification (30 cycles):<br>Denaturation: 30 sec 95°C<br>Annealing: 30 sec 65°C<br>Extension: 3 min 72°C<br>Final extension: 5 min 72°C |
| Pf6-Rev-Schmidt2023<br>Pf6-Fwd1-Schmidt2023<br>Pf6-Fwd2-Schmidt2023 | attR of Pf6 (if integrated) or 5' – 3' sites of Pf6 (bacterial genome when phage is lost) | EconoTaq PLUS GREEN 1X Master Mix, 25 pmol primers. Template: 1 µl lysate or 50 ng phage ssDNA | Initial denaturation: 2 min 95°C.<br>Amplification (30 cycles):<br>Denaturation: 30 sec 95°C<br>Annealing: 30 sec 64°C<br>Extension: 2 min 72°C<br>Final extension: 5 min 72°C |
| Fw-Pf6-RF-Saumel2024<br>Rv-Pf6-RF-Saumel2024 | Pf6 RF form | EconoTaq PLUS GREEN 1X Master Mix, 50 pmol primers. Template: 1 µl lysate or 30 ng phage ssDNA | Initial denaturation: 2 min 95°C.<br>Amplification (35 cycles):<br>Denaturation: 30 sec 95°C<br>Annealing: 30 sec 62°C<br>Extension: 30 sec 72°C<br>Final extension: 5 min 72°C |
| Pf4_mut_FW<br>Pf4_mut_RV | Pf4 phage | EconoTaq PLUS GREEN 1X Master Mix, | Initial denaturation: 2 min 95°C.<br>Amplification (30 cycles): |

| | | | |
|---|---|---|---|
| | | 25 pmol primers.<br>Template: 1 µl lysate or 30 ng phage ssDNA | Denaturation: 30 sec 95°C<br>Annealing: 30 sec 64°C<br>Extension: 1 min 72°C<br>Final extension: 5 min 72°C |
| M13_pIII_FW<br>M13_pIII_RV | gIII gene, M13 phage | EconoTaq PLUS GREEN 1X Master Mix, 50 pmol primers.<br>Template: 1 µl lysate or 30 ng phage ssDNA | Initial denaturation: 2 min 95°C.<br>Amplification (30 cycles):<br>Denaturation: 30 sec 95°C<br>Annealing: 30 sec 63°C<br>Extension: 1 min 72°C<br>Final extension: 5 min 72°C |
| K12IS-L<br>K12-R | K-12 rfb locus | EconoTaq PLUS GREEN 1X Master Mix, 25 pmol primers.<br>Template: 1 µl lysate or 30 ng phage ssDNA | Initial denaturation: 2 min 95°C.<br>Amplification (30 cycles):<br>Denaturation: 30 sec 95°C<br>Annealing: 30 sec 63.5°C<br>Extension: 1 min 30 sec 72°C<br>Final extension: 5 min 72°C |

**Supplementary Table III. Purification methods for phage DNA.**

| Year | Citation | Method | Method summary | Sample/ Phage | DNA classification dsDNA/ssDNA/linear-circular | Yield | Quality | Downstream applications |
|---|---|---|---|---|---|---|---|---|
| 2014 | [6] | DNA Clean & Concentrator-5 kit | Dissolved plaque treated with DNase I, proteinase K, SDS and heat. Silica-based spin column; protocols and buffers from the manufacturer. | Φ29, P113g, T4 | Φ29: 19 kb linear dsDNA with ends covalently connected to gp3.<br>P113g: 30.8 kb linear dsDNA<br>T4: 169 kb linear dsDNA | ND | ND | Sequencing by Illumina |
| 2018 | [7] | DNeasy Blood & Tissue Kit | Concentrated phages in SM + gelatin buffer recovered from double agar plates. 450 µl out of 5 ml per plate treated with DNase I, RNase A, proteinase K and heat. Lysate, AL buffer, and EtOH mixed 1:1:1 (v/v) and applied to a silica-based column. | 16 phages (SU) against Streptococcus uberis; Salmonella (S112) and Escherichia coli (EC11 and EC80) | dsDNA | 136.8 – 8,620.5 ng | - A260/280: 84% samples >1.80<br>- No degradation or RNA contamination in agarose gel | Sequencing by Illumina |
| 2021 | [8,9] | Phage DNA Isolation Kit (Norgen Biotek) | 1 ml of chloroform-treated SN. Heat, chemical and proteinase K lysis. Column-based protocol. | T7 phage | 40 kb linear dsDNA | 3-15 µg | Some smear in agarose gel | NGS Illumina |
| 2021 | [10] | Acetone - kit | Phage SN treated with DNase I and RNase I, phages precipitated by acetone and purified with the PureLink Viral RNA/DNA Mini Kit. Proteinase K and lysis buffer treatment, EtOH addition, and loading onto a silica column. | - E. faecalis phage<br>- Staphylococcus aureus phage (50 kb) | dsDNA | 5 µg/ml SN | - $A_{260/280}$: 1.99<br>- Restriction<br>- No degradation | None |
| | | Acetone – Iodide salts | Phage SN treated with DNase I and RNase I, phages | | | 5 µg/ml SN | - $A_{260/280}$: 1.99<br>- | Illumina HiSeq 2500 |

| | | | | | | | | |
|---|---|---|---|---|---|---|---|---|
| | | | precipitated by acetone. Mixed 1:1 (v/v) with 3 M iodide salt and loaded onto a silica column. | | | | Restriction<br>- No degradation | |
| | | TCA - kit | Same as acetone, but phages precipitated with TCA | | | 3.38 µg/ml SN | $A_{260/280}$: 1.99 | None |
| | | TCA – Iodide salts | | | | 3.13 µg/ml SN | $A_{260/280}$: 1.94 | |
| | | PEG - kit | Same as acetone, but phages precipitated by PEG/NaCl | | | 4.63 µg/ml SN | $A_{260/280}$: 1.96 | |
| | | PEG – Iodide salts | | | | 4.44 µg/ml SN | $A_{260/280}$: 1.93 | |
| | | UCF - kit | Same as acetone, but phages collected by ultracentrifugation | | | 5 µg/ml SN | $A_{260/280}$: 2.25 | |
| | | UCF – Iodide salts | | | | 4.63 µg/ml SN | $A_{260/280}$: 2.06 | |
| 2022 | 11 | NEB Monarch kit | SN filtered, PEG-precipitated, and purified with the NEB Monarch kit | E. coli T4 | linear dsDNA 169 kb | 176 ng/ml SN | - $A_{260/280}$: 1.90<br>- $A_{260/230}$: 1.79<br>- Restriction | Chemical analyses (HPLC and LC–MS/MS) |
| | | | | B. subtilis SPO1 | linear dsDNA 146 kb | 604 ng/ml SN | - $A_{260/280}$: 2.18<br>- $A_{260/230}$: 1.99<br>- Restriction | |
| 2022 | 12 | QIAamp MinElute Virus Spin Kit | VLPs extracted from feces in SM buffer, concentrated, washed, treated with DNase, and purified with the kit | Virus-like particles (VLPs) | dsDNA | 160 ng / 250 mg feces | ND | NGS Illumina |
| 2022 | 13 | GuHCl/ EtOH-Glass membrane | Filtered SN, PEG/NaCl, CsCl gradient, DNase and RNase. Phage lysis (proteinase K, heat, lysis buffer), EtOH and column procedure | G. stearothermophilus TP-84 | dsDNA 47.7 kb | ND: 7.1 µg on average from unknown SN volume | - $A_{260/280}$: 1.92<br>- $A_{260/230}$: 1.91<br>- Restriction | None |
| | | GuHCl/ | | | | ND: | - $A_{260/280}$: | |

| | | | | | | | | |
|---|---|---|---|---|---|---|---|---|
| | | EtOH-Silica column | | | | 5.2 µg on average from unknown SN volume | 2.01<br>- $A_{260/230}$: 2.36<br>- Restriction | |
| 2022 | [14] | Monarch HMW DNA Extraction Kit | Phage SN PEG-precipitated, DNase and RNase treated, phage lysis (proteinase K, lysis buffer, heat and agitation), followed by the kit instructions (capture on glass beads) | X. oryzae XP12 | linear dsDNA 63.7 kb | 0.9-4.5 µg/ml SN | - $A_{260/280}$: 1.72<br>- $A_{260/230}$: 2.05<br>- Pulsed field gel electrophoresis: no smearing | Nucleotide analyses by UHPLC |
| | | | | B. subtilis SP8 | linear dsDNA 138.7 kb | 0.64 – 3.2 µg/ml SN | - $A_{260/280}$: 1.99<br>- $A_{260/230}$: 2.09<br>- Pulsed field gel electrophoresis: minimal smearing | |
| | | | | E. coli CBA120 | linear dsDNA 157.3 kb | 1.8 - 9 µg/ml SN | - $A_{260/280}$: 2.30<br>- $A_{260/230}$: 1.90<br>- Pulsed field gel electrophoresis: minimal smearing | |
| | | | | E. coli T4 | linear dsDNA 168.9 kb | 1.62 – 8.1 µg/ml SN | - $A_{260/280}$: 1.77<br>- $A_{260/230}$: 1.97<br>- Pulsed field gel electrophoresis: minimal smearing | |
| | | | | B. megateri | linear dsDNA 497.5 kb | 0.48 – 2.4 | - $A_{260/280}$: 1.80 | |

| | | | | | | | | |
|---|---|---|---|---|---|---|---|---|
| | | | | um G (Priestia megaterium) | | μg/ml SN | - $A_{260/230}$: 1.88<br>- Pulsed field gel electrophoresis: extensive smearing | |
| 2024, 2009 | [15,16] | Promega Wizard DNA Clean-Up kit | Phage SN treated with DNase I and RNase I, phages precipitated by PEG, lysed with resin in guanidinium thiocyanate. Resin composition is proprietary | Lytic phages | dsDNA | 0.1 – 1 μg/ml SN | ND | None |
| 2010 | [17] | Promega Wizard DNA Clean-Up kit | Unknown | M13mp18 | ssDNA | ND | ND | None |
| 2019 | [18] | EndoFree Plasmid Maxi Kit | Filtered PEG-precipitated SN, phage lysis (proteinase K, P2 buffer, heat), P3 buffer, filtering, EtOH addition and anion-exchange resin | M13cp helper plasmid / mini-phagemid | ssDNA | 2.0 μg/ml SN | 88% purity on agarose gel | DNA origami assembly |
| 2019 | [19] | E.Z.N.A.® M13 DNA Mini Kit | Phage precipitated from SN, bound to HiBind matrix and lysed, washed and eluted | M13 | ssDNA | 1 – 3.3 μg/ml SN | ND | None |
| 2020 | [20] | Chemical dissociation – QIAGEN MiniPrep Kit | Neat sputum diluted 1:1 with PBS, DNase and DTT, heat-treated and purified according to the kit. | Pf phages and Pseudomonas aeruginosa | Phage ssDNA and genomic dsDNA from P. aeruginosa | From 50 μl neat sputum: 2 μg | ND | qPCR |
| | | Mechanical dissociation - QIAamp DNA Mini Kit | Neat sputum diluted 1:1 with AL buffer, supplemented with proteinase K, and disrupted by bead beating. | | | From 200 μl neat sputum: 20 μg | | |
| 2021 | [21] | Nucleo Spin | PEG-precipitated phage SN, phage | M13mp18 | ssDNA | 2 μg/ml | No smear | DNA origami |

| | | | | | | | | |
|---|---|---|---|---|---|---|---|---|
| | | Gel and PCR Clean-up (Macherey-Nagel) / Exprep plasmid SV mini (GeneAll Biotechnology) | lysis (P2 buffer, heat), P3 buffer, addition of binding buffer to SN (chaotropic salts) and application to a silica column | | | SN | on agarose gel | assembly |
| | | | | M13mp18 mut62 | | 12 µg/ml SN | | |

**ND:** Not disclosed; SN: Supernatant

**Supplementary Table IV. Summary of prediction results for oligonucleotide dimerization by the DINAMelt server.**

| Parameter | ***KpnI* oligo** | ***EcoRI* oligo** | ***NdeI* oligo** | ***SmaI* oligo** |
|---|---|---|---|---|
| ΔG (kcal/mol) | -8.2 | -9.7 | -12.2 | -12.7 |
| ΔH (kcal/mol) | -59.0 | -74.8 | -79.0 | -123.2 |
| ΔS (e.u.) | -163.7 | -209.8 | -215.5 | -356.2 |
| Tm (°C) | 43.0 | 48.3 | 58.3 | 51.9 |

Calculation conditions: 37°C, [$Na^+$] 50 mM, [$Mg^{2+}$] 10 mM.

**Supplementary Table V. Summary of ssDNA yields of M13 phage from literature.**

| Year | Citation | Phage system | Purification method | Yield | Quality Control |
|---|---|---|---|---|---|
| 1990 | [22] | M13mp18 / TG1 in 2XTY / 96 well plate / batch | PEG-NaCl precipitation / phage lysis (SDS-heat) / EtOH-NaAc precipitation / wash / ND quantification | 6.67 μg/ml SN | Sequence |
| 1992 | [23] | M13mp18 / Cells in 2xTy / Tubes / batch | Phage lysis (SDS-heat) followed by addition of PEG-NaCl and biotinylated probe complementary to M13 and streptavidin coated magnetic beads / wash / elution / ND quantification | 1.5 μg/ml SN | Sequence |
| 1992 | [24] | M13mp18 / DH5αF' in 2xYT / Deep well plate / batch | PEG-NaCl + magnetic beads precipitation / phage lysis / wash-elution / agarose gel quantification | 2.5 μg/ml SN | Sequence |
| | | | Phenol / agarose gel quantification | 4 μg/ml SN | |
| 1993 | [25] | M13 / DH5αF' in 2xYT / 96 deep well plate / batch | PEG-NaCl precipitation / Phage lysis (KI-KDS-heat) / Agarose gel quantification | 10-20 μg/ml SN | Sequence |
| 1993 | [26] | M13 vector / M13 E. coli in 2XTY / Tubes / batch | PEG-NaCl precipitation / Phenol-chloroform-ethanol precipitation / Agarose gel quantification | 5-10 μg/ml SN | Sequence |
| 1994 | [27] | M13mp19 / DH5αF' in 2xYT / 48 well plate / batch | Phage lysis (NaClO4) / Capture on glass fiber-filter plate / wash-elution / Agarose gel quantification | 0.86 – 1.71 μg/ml SN / 5-7 x $10^{11}$ PFU | Sequence |
| 1996 | [28] | M13mp19 / TG1 in 2XYT / Tubes / batch | PEG-NaCl precipitation / Capture on glass fiber-filter plate / wash-elution / Agarose gel quantification | 3-5 μg/ml SN | Sequence Bacterial RNA contaminant |
| | | | Phage lysis (NaClO4) / Capture on glass fiber-filter plate / wash-elution / Agarose gel quantification | | |
| 1996 | [29] | M13 / JM109 in 2XTY / batch | Phage lysis (SDS-heat) followed by SDS removal (NaOAc-PEG/centrifugation) and addition of biotinylated probe complementary to M13 and streptavidin coated magnetic beads to SN /wash / elution / Agarose gel quantification | 1.5 μg/ml SN | Sequence <5% host DNA contaminant |
| 1996 | [30] | M13mp18 / TG1 in LB / 15 ml Tubes / batch | PEG-NaCl precipitation / Phenol / NaAc-ethanol precipitation / OD quantification | 10 – 15 μg/ml SN / 4.6 – 8 x $10^{11}$ | Sequence |

| | | | | | |
|---|---|---|---|---|---|
| | | | | PFU | |
| 2015 | [31] | Phagemid pSB4434 / plasmid-encoded helper phage / S2188 in rich media / batch | PEG-NaCl precipitation / Phenol-chloroform extraction / ethanol precipitation / OD quantification | 0.4 µg/ml SN | dsDNA contamination |
| 2015 | [32] | M13mp18 / XL1-blue MRF' Kan in defined medium with feed solution / fed-batch | PEG-NaCl precipitation / phage lysis (P2 and P3 buffers Qiagen) / ethanol precipitation / OD quantification | 370 – 410 µg/ml SN / 1.6 x $10^{14}$ PFU / 67.7 mg cell dry weight | No additional DNA on agarose gel |
| 2017 | [33] | M13mp18 / XL1-blue MRF' Kan in defined medium with feed solution / fed-batch | PEG-NaCl precipitation / phage lysis (P2 and P3 buffers Qiagen) / ethanol precipitation / Fluorescence quantification | 590 µg/ml SN / $10^{14}$ PFU | ND |
| | | | | 320 µg/ml SN / $10^{14}$ PFU / 65 mg cell dry weight | ND |
| 2017 | [34] | M13mp18 / JM109 in defined medium and glucose feed / continuous operation | PEG-NaCl precipitation / phage lysis (P2 and P3 buffers Qiagen) / ethanol precipitation / Fluorescence quantification | 10.6 µg/ml SN / 7 mg cell dry weight | ND |
| | | | | 6 µg/ml SN / 9 mg cell dry weight | |
| | | | | 25 µg/ml SN / 28 mg cell dry weight | |
| 2019 | [18] | M13cp helper plasmid / mini-phagemid / SS320 in 2xYT / shake flask / batch | Phage lysis (Qiagen P1-proteinase K-RNAse -Qiagen P2-heat -Qiagen P3-centrifugation) / ethanol addition / Qiagen spin column / OD quantification | 2.3 µg/ml SN | 88% purity on agarose gel |
| | | M13cp helper plasmid / mini-phagemid / SS320 in 2xYT / fermenter/ batch | PEG-NaCl / Qiagen EndoFree Maxiprep kit with phage lysis (proteinase K-heat), endotoxin removal and ssDNA column purification (EtOH) / OD | 2.0 µg/ml SN | No additional DNA on agarose gel |

| | | | | | |
|---|---|---|---|---|---|
| | | | quantification | | |
| 2021 | [35] | Phagemid / VCSM13 helper phage / JM109 in supplemented 2xYT / bottle / batch | PEG-NaCl precipitation / phage lysis (NaOH-SDS - KOAc -centrifugation) / ethanol precipitation / weighing of dried material for quantification | 3.13 µg/ml SN | Agarose gel |
| 2021 | [21] | M13mp18 / XL1-Blue in 2XM9 medium / shake flask / batch | PEG-NaCl precipitation / phage lysis (NaOH-SDS-heat - NaAc-centrifugation) / column purification (Exprep plasmid SV mini) / OD quantification | 2 µg/ml SN | Agarose gel |
| | | M13mp18 mut62 / XL1-Blue in 2XM9 medium / shake flask / batch | | 12 µg/ml SN | |
| | | M13mp18 / XL1-Blue in 2XM9 medium and feed/ stirred tank reactor / fed-batch | | 56.14 µg/ml SN / 1.27 x $10^{15}$ PFU / 21.6 mg cell dry weight | |
| | | M13mp18mut62 / XL1-Blue in 2XM9 medium and feed/ stirred tank reactor / fed-batch | | 268.41 µg/ml SN / 9.7 x $10^{15}$ PFU / 21.6 mg cell dry weight | |
| 2022 | [36] | Phage-encapsulated phagemid / Helper plasmid / JM109 in defined medium and feed / stirred tank bioreactor / fed-batch 37 °C | PEG-NaCl precipitation / phage lysis (Qiagen P2-P3-centrifugation) / ethanol precipitation / OD quantification | 11 µg/ml SN / 100 mg cell dry weight | 28% purity on agarose gel |
| | | Phage-encapsulated phagemid / Helper plasmid / JM109 in defined medium / shake flask / batch 30 °C | | 4.15 µg/ml SN | ND |
| | | Phage-encapsulated phagemid / Helper plasmid / JM109 in defined medium / shake flask / batch 37 °C | | 1.36 µg/ml SN | ND |
| | | Phage-encapsulated phagemid / Helper plasmid / JM109 in defined medium and feed / stirred tank bioreactor / fed-batch | | 83 µg/ml SN / 95 mg cell dry weight | 61% purity on agarose gel |

| | | | | | |
|---|---|---|---|---|---|
| | | 30 °C | | | |
| 2024 | [37] | Phage-encapsulated phagemid / Helper plasmid / JM109 in defined medium and feed / 2.5 L bioreactor / fed-batch | PEG-NaCl precipitation / phage lysis (Qiagen P2-P3-centrifugation) / ethanol precipitation / OD quantification | 70.84 µg/ml SN / 80 mg cell dry weight | Agarose gel |
| | | | | 127.22 µg/ml SN / 49.58 mg cell dry weight | |
| | | Phage-encapsulated phagemid / Helper plasmid / JM109 in defined medium and feed / 25 L bioreactor / fed-batch | | 191.08 µg/ml SN / 50.6 mg cell dry weight | |
| | | | | 159.10 µg/ml SN / 45.5 mg cell dry weight | |
| | | Phage-encapsulated phagemid / Helper plasmid / JM109 in defined medium and feed / 1000 L bioreactor / fed-batch | | 55.34 µg/ml SN / 48 mg cell dry weight | |
| 2024 | [38] | Phagemid / Helper strain TG1 in LB medium / shake flasks / batch | PEG-NaCl precipitation / phage lysis (Qiagen P2-P3-centrifugation) / ethanol precipitation / OD quantification | 0.25 µg/ml SN | 95–98% purity Agarose gel |
| 2024 | [39] | M13mp18 / ER2738 in 2xYT / shake flask / batch | PEG-NaCl precipitation / phage lysis (P2 and P3 buffers Qiagen) / ethanol precipitation / Ion-Pair Reverse Phase Chromatography quantification | 12.6 µg/ml SN | IP-RP chromatography |

**ND:** Not disclosed; **SN:** Supernatant; **OD:** Optical density.

### Supplementary Figures

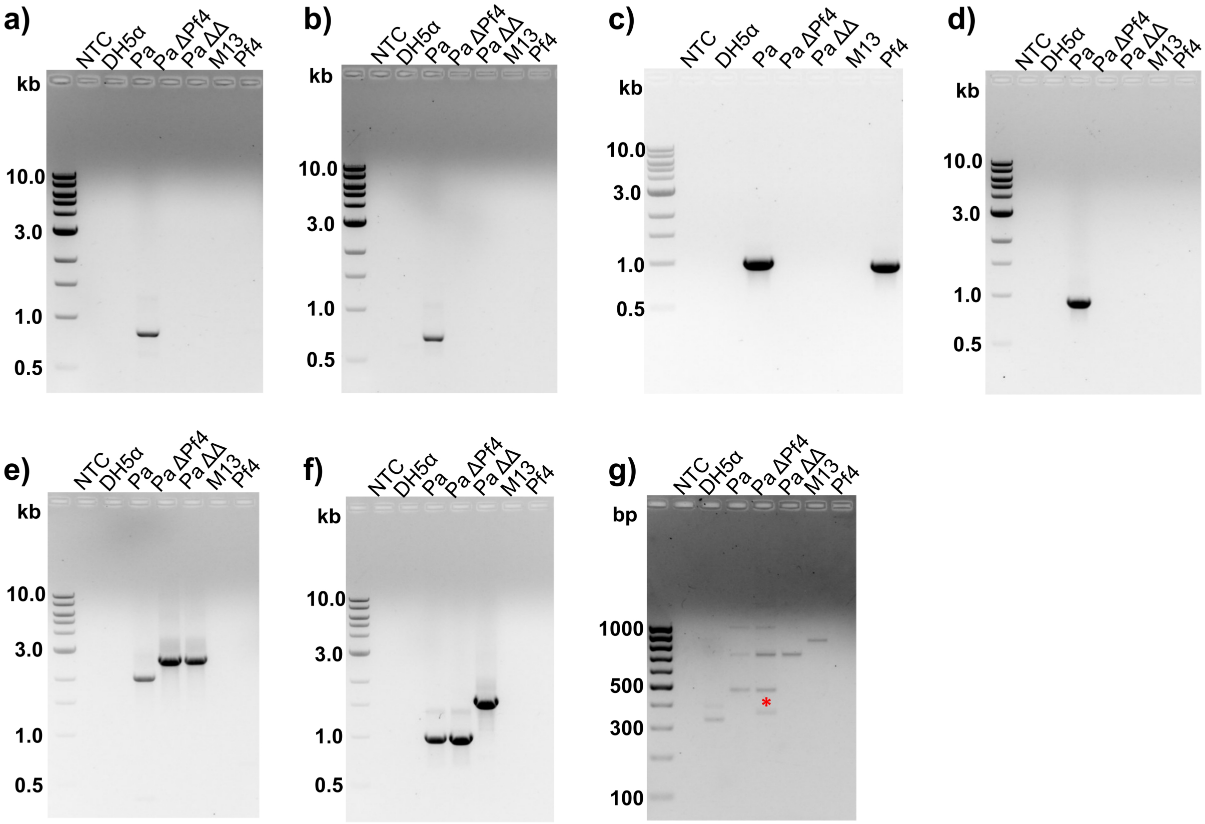


**Supplementary Figure 1.** PCR validation of *Pseudomonas* strains MPAO1, MPAO1 ΔPf4 and MPAO1 ΔPf4 ΔPf6. **a)** Detection of the Pf4 replicative form (RF) with primers Pf4F-Webb2004/Pf4R-Webb2004. 839 bp amplicon. **b)** Amplification of the bacteria–prophage junction at the attL site with primers GendelF-Rice2009/GendelR-Rice2009. 806 bp amplicon. **c)** Amplification of two Pf4-specific genes with primers IntF4-F-Gavric2022/IntF4-R-Gavric2022. 1001 bp amplicon. **d)** Amplification of the bacteria–prophage junction at the attR site with primers Pf4/PA14-F-Gavric2022/Pf4/PA14-R-Gavric2022. 943 bp amplicon. **e)** Multiplex PCR with primers Pf4-Rev-Schmidt2023/Pf4-Fwd1- Schmidt2023/Pf4-Fwd2- Schmidt2023 giving a 2020 bp amplicon when the Pf4 prophage is present and a 2570 bp amplicon when it is lost. **f)** Multiplex PCR with primers Pf6-Rev- Schmidt2023/Pf6-Fwd1- Schmidt2023/Pf6-Fwd2-Schmidt2023 giving a 1016 bp amplicon when the Pf6 prophage is present and a 1572 bp amplicon when it is lost. **g)** Detection of the Pf6 RF with primers Fw-Pf6-RF-Saumel2024/Rv-Pf6-RF-Saumel2024. 353 bp amplicon indicated by a red asterisk above the band. NTC: non-template control; DH5α: Cell lysate from *E. coli* DH5α strain; Pa: Cell lysate from *Pseudomonas aeruginosa* MPAO1 strain; Pa ΔPf4: Cell lysate from *Pseudomonas aeruginosa* MPAO1 ΔPf4 strain; Pa ΔΔ: Cell lysate from *Pseudomonas aeruginosa* MPAO1 ΔPf4 ΔPf6 strain; M13: Purified ssDNA from the *E. coli* phage M13KO7; Pf4: Purified ssDNA from *P. aeruginosa* phage Pf4. kb: kilobases. A 1 kb DNA ladder was used in panels **a-f** and a 100 bp ladder in panel **g**.

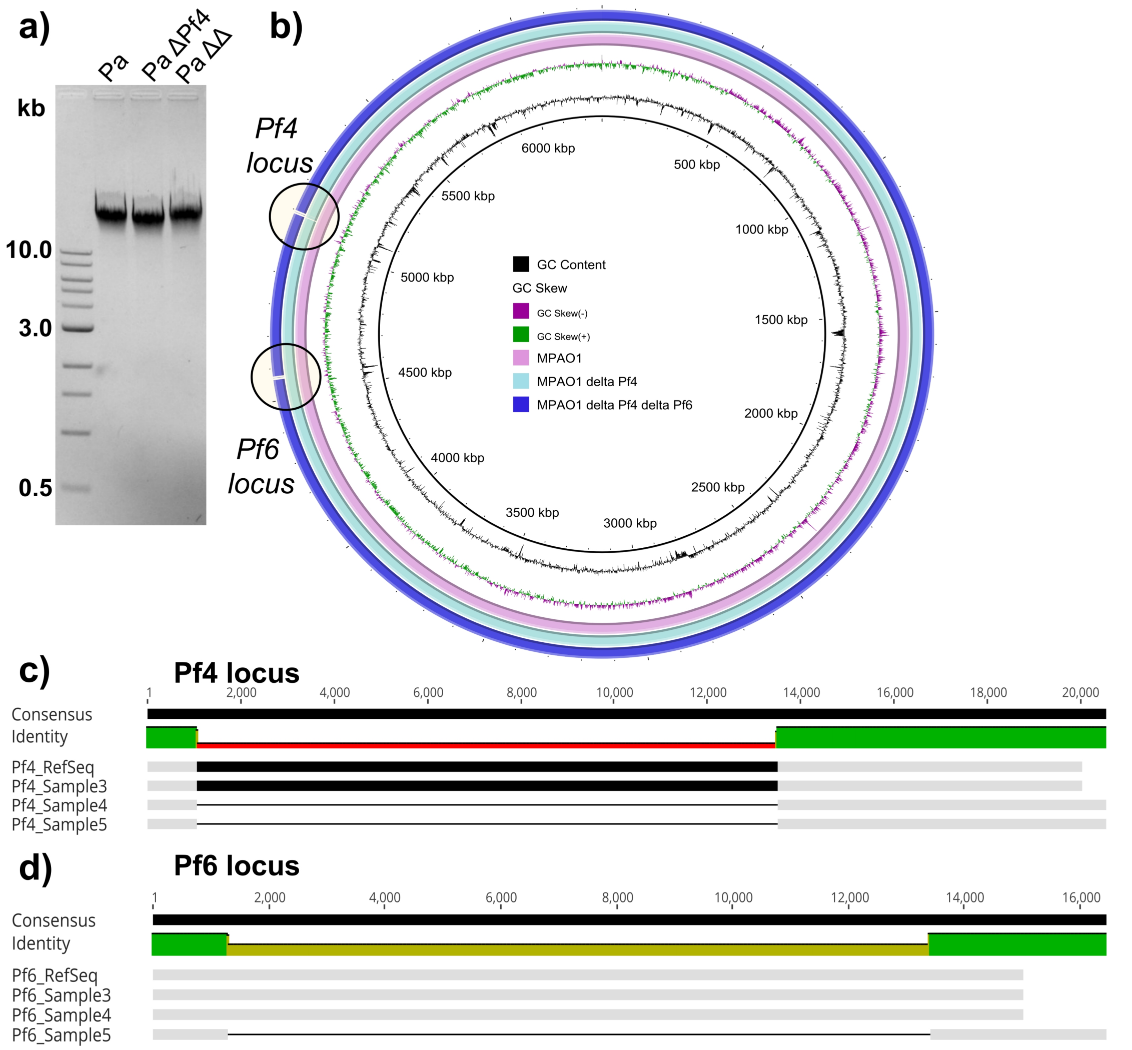


**Supplementary Figure 2.** Purification, sequencing, and analysis of gDNA from *Pseudomonas aeruginosa* strains. **a)** Quality control of the purity and integrity of purified gDNA by visualization on a 1% agarose gel alongside a 1 kb DNA ladder. kb: kilobases. Pa: *Pseudomonas aeruginosa* MPAO1 strain; Pa ΔPf4: *Pseudomonas aeruginosa* MPAO1 ΔPf4 strain; Pa ΔΔ: *Pseudomonas aeruginosa* MPAO1 ΔPf4 ΔPf6 strain. **b)** Whole-genome alignment in the BLAST Ring Image Generator (BRIG), highlighting the Pf4 and Pf6 prophage loci. **c)** Multiple alignment of the Pf4 and Pf6 prophages across 20 kb in Geneious Prime.

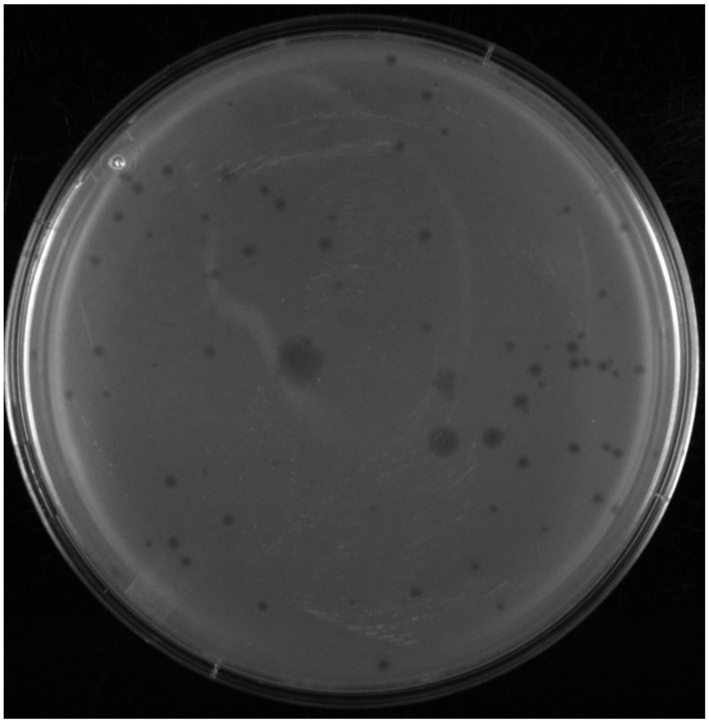

**Supplementary Figure 3.** Petri dish showing individual Pf4 plaques isolated from the laboratory Pf4 phage bank. A 100 µl volume of a $10^7$ dilution of clarified supernatant from MPAO1 ΔPf4 ΔPf6 strain infected with Pf4 phage was mixed with 20 ml of LB Miller 0.5% agar, poured into a Petri dish, and incubated for 14–16 h at 37°C.

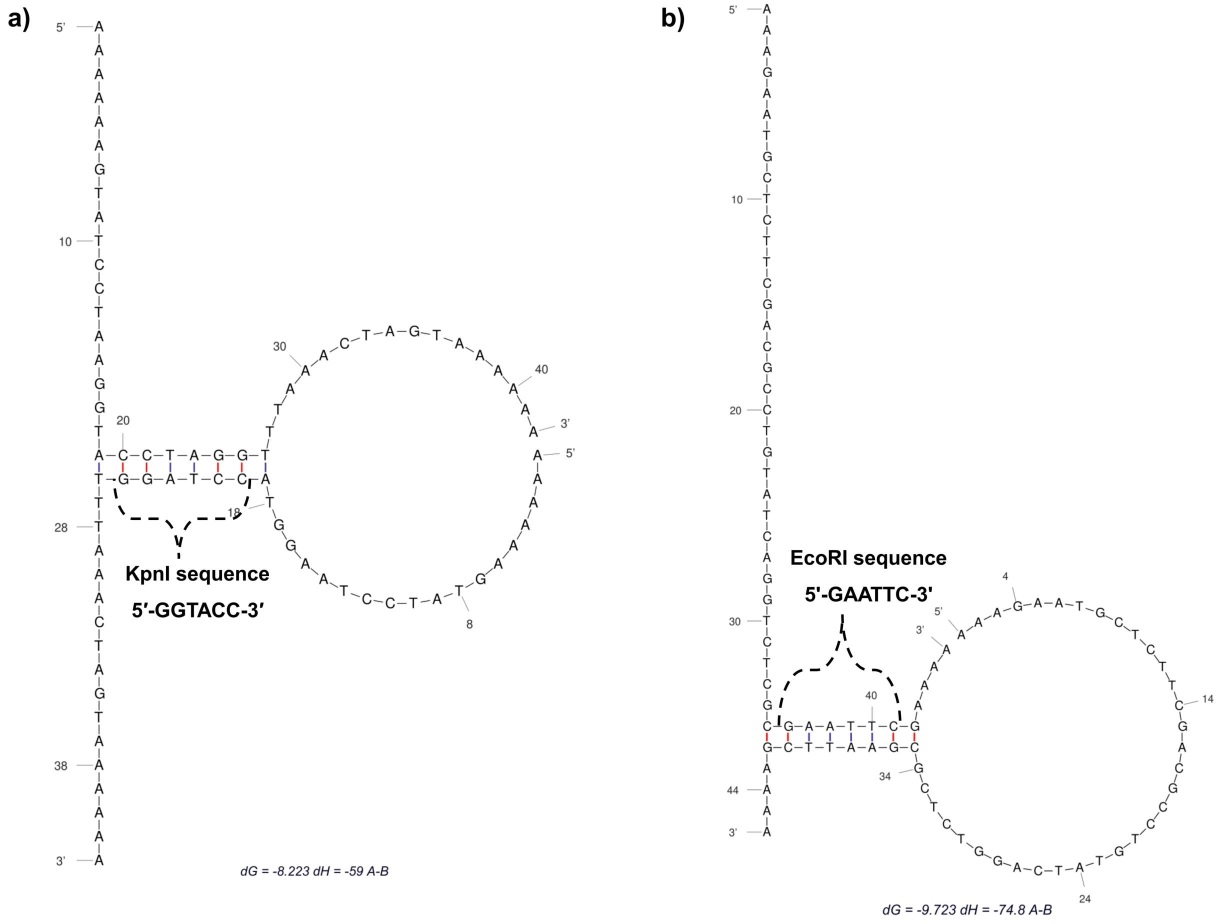


**Supplementary Figure 4.** Calculation of homodimer formation for *KpnI* and *EcoRI* REBASE oligonucleotides. Homodimer formation was predicted with the DINAMelt server, using the two-state melting (hybridization) application for the oligonucleotides in the REBASE database for ssDNA cleavage by *KpnI* (**a**) and *EcoRI* (**b**). Calculations were performed for DNA at 37 °C, approximating the ionic conditions of rCutSmart buffer. All other parameters were left at their default values.

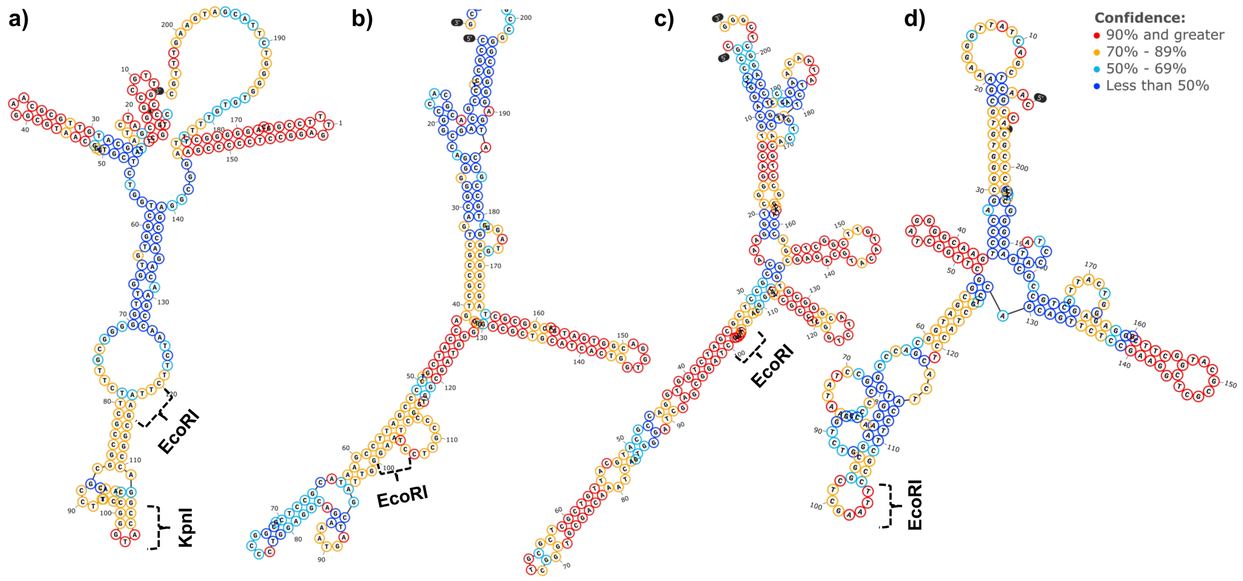


**Supplementary Figure 5.** Calculation of local structures in the Pf4 ssDNA genome using SnapGene with ViennaRNA DNA options. Local folding around the *KpnI* site (**a**) and *EcoRI* sites (**a-d**) is shown with the recognition sites marked. For paired bases, the color denotes the probability that the bases are paired. For unpaired regions, the color denotes the probability of being unpaired.

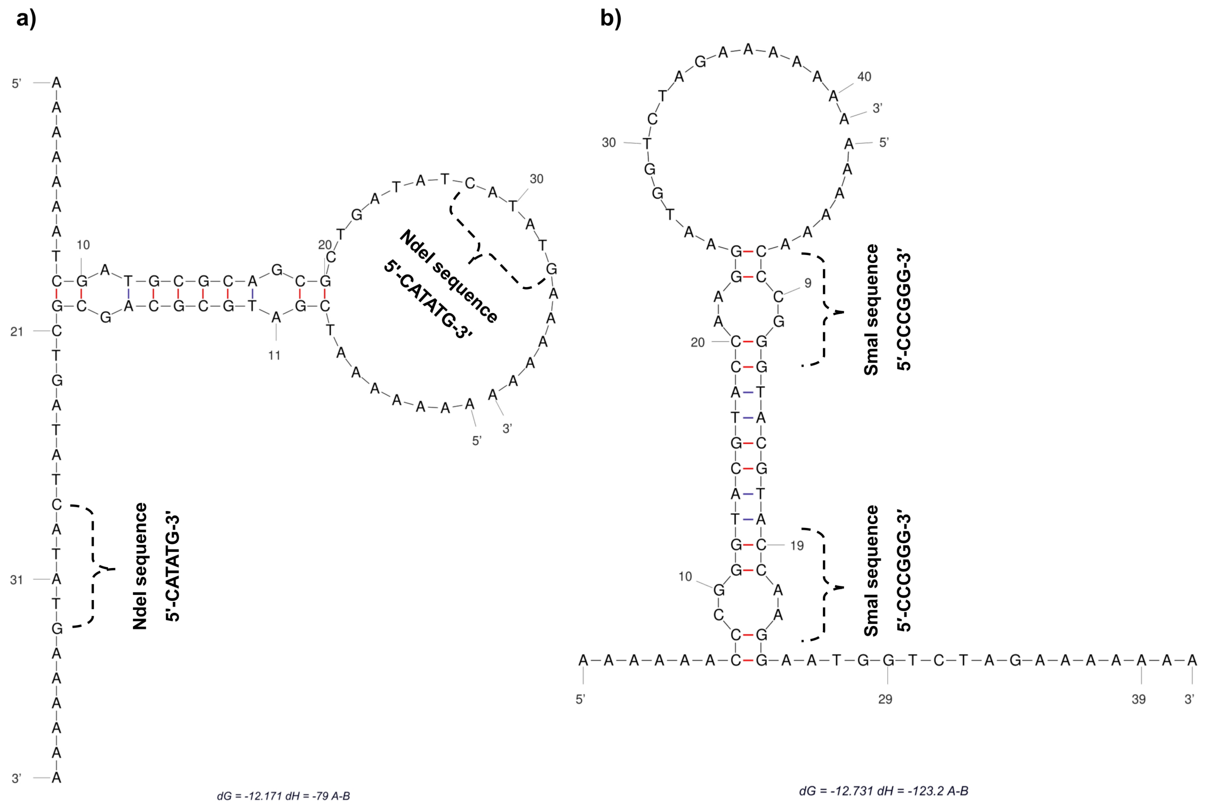


**Supplementary Figure 6.** Calculation of homodimer formation for *NdeI* and *SmaI* REBASE oligonucleotides. Homodimer formation was predicted with the DINAMelt server, using the two-state melting (hybridization) application for the oligonucleotides in the REBASE database for ssDNA cleavage by *NdeI* (**a**) and *SmaI* (**b**). Calculations were performed for DNA at 37 °C, approximating the ionic conditions of rCutSmart buffer. All other parameters were left at their default values.

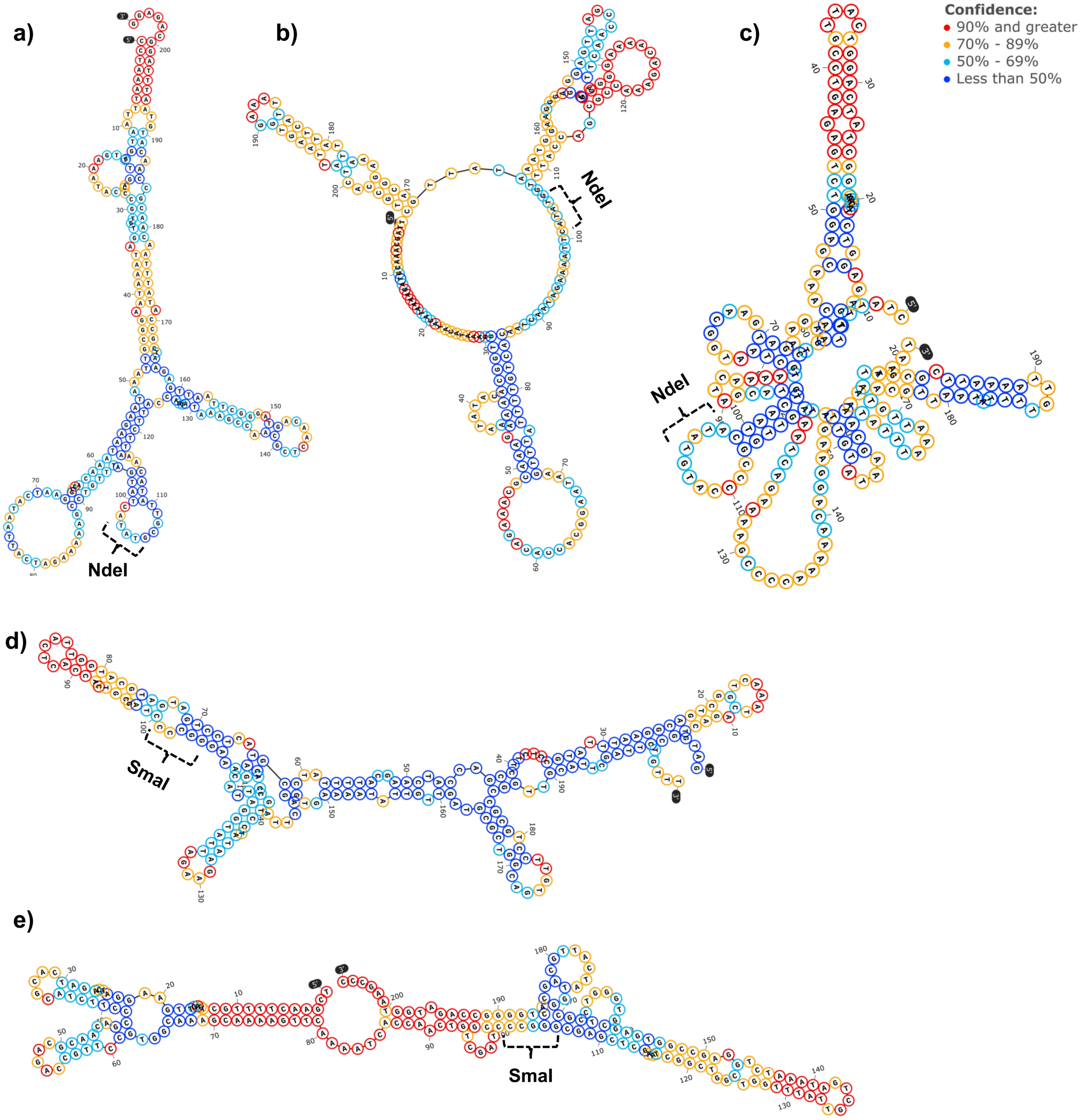


**Supplementary Figure 7.** Calculation of local structures in the M13KO7 ssDNA genome using SnapGene with ViennaRNA DNA options. Local folding around *NdeI* sites (**a-c**) and *SmaI* sites (**d-e**) is shown with the recognition sites marked. For paired bases, the color denotes the probability that the bases are paired. For unpaired regions, the color denotes the probability of being unpaired.
